# The costimulatory domain influences CD19 CAR-T cell resistance development in B-cell malignancies

**DOI:** 10.1101/2025.02.28.640707

**Authors:** Marta Krawczyk, Narcis Fernandez-Fuentes, Klaudyna Fidyt, Tomasz Winiarski, Monika Pepek, Agnieszka Graczyk-Jarzynka, Jacinta Davis, Pablo Bousquets-Muñoz, Xose S. Puente, Pablo Menendez, Emmanuelle Benard, Sébastien Wälchli, Andrei Thomas-Tikhonenko, Magdalena Winiarska

## Abstract

CD19-CAR-T-cells emerge as a major therapeutic option for relapsed/refractory B-cell-derived malignancies, however approximately half of patients eventually relapse. To identify resistance-driving factors, we repeatedly exposed B-cell lymphoma/B-cell acute lymphoblastic leukemia to 4-1BB/CD28-based CD19-CAR-T-cells *in vitro*. Generated models revealed costimulatory domain-dependent differences in CD19 loss. While CD19-4-1BB-CAR-T-cells induced combination epitope/total CD19 protein loss, CD19-CD28-CAR-T-cells did not drive antigen-escape. Consistent with observations in patients relapsing after CD19-4-1BB-CAR-T-cells, we identified CD19 frameshift/missense mutations affecting residues critical for FMC63 epitope recognition. Mathematical simulations revealed that differences between CD19-4-1BB- and CD19-CD28-CAR-T-cells activity against low-antigen-expressing tumor contribute to heterogeneous therapeutic responses. By integrating *in vitro* and *in silico* data, we propose a biological scenario where CD19-4-1BB-CAR-T-cells fail to eliminate low-antigen tumor cells, fostering CAR-resistance.

These findings offer mechanistic insight into the observed clinical differences between axi-cel (CD28-based) and tisa-cel (4-1BB-based)-treated B-cell lymphoma patients and advance our understanding on CAR-T resistance. Furthermore, we underscore the need for specific FMC63 epitope detection to deliver information on antigen levels accessible for CD19-CAR-T-cells.

**Visual abstract:** 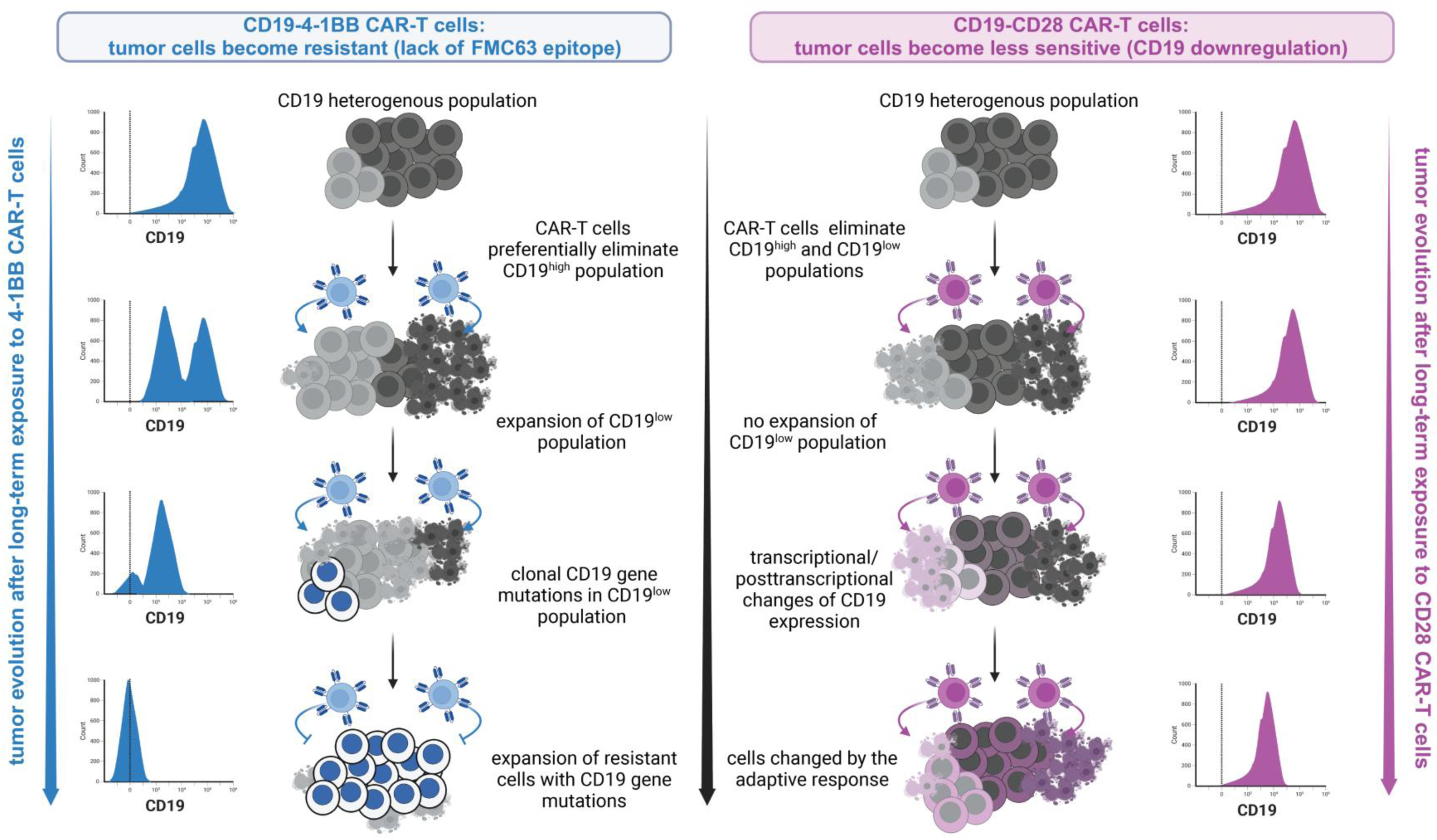

## Introduction

In recent years, phenomenal progress has been witnessed in hemato-oncology related to the implementation of cancer immunotherapies. These therapeutic strategies involve the body’s immune system to recognize, attack and kill tumor cells. The development of adoptive cell therapy with CAR-redirected immune cells against CD19 antigen was a major breakthrough in the hemato-oncology field. Since 2017, four different anti-CD19 CAR-T products based on FMC63 scFv were approved for the treatment of malignancies derived from abnormal B cells at different stages of their differentiation, mostly relapsed/refractory (r/r) diffused large B-cell lymphoma (DLBCL) and r/r B-cell acute lymphoblastic leukemia (B-ALL). These approved products incorporate two different costimulatory domains, namely 4-1BB (Kymriah, Tisagenlecleucel (tisa-cel) and Breyanzi, Lisocabtagene Maraleucel (liso-cel)) and CD28 (Yescarta, Axicabtagene Ciloleucel (axi-cel) and Tecartus, Brexucabtagene Autoleucel (brexu-cel)). The approval of these products was based on phenomenal results in initial clinical trials where complete remission (CR) rates reached 40-79 % [1–7]. Similar efficacy of these therapies were confirmed by the long-term observation studies with the median follow-up range from 24 up to 61 months in lymphoma [8–15] and 13-57.6 months in B-ALL patients [16–25]. However, despite the phenomenal success of CD19 CAR-T therapy limitations and side effects are still observed and need to be further explored.

In particular, the clinical outcomes indicate that a significant part of patients with high-grade lymphomas and B-ALL exhibit primary or acquired resistance. The most frequently reported causes of resistance to CD19 CAR-T therapy include aberrations of malignant cells, with the most commonly described CD19 antigen loss [26,27]. The mechanisms responsible for antigen loss include primarily mutations of CD19 gene and alternative splicing of the CD19 transcript. The alternative splice variants lacking exon 2 or exons 5-6 as well as mutations affecting exons 2-5 were detected in samples from relapsed B-ALL patients after CD19 CAR-T treatment. All mentioned aberrations led to the expression of a non-synonymous variant or truncated protein lacking the epitope recognized by CD19 CAR-T cells and thus contributing to the development of resistance [28,29]. Based on the FMC63 epitope mapping, it is known that exon 3 and 4 encode the residues (W159, R163, K220, P222) crucial for FMC63 formation and proper recognition by the FMC63-based CD19 CAR-T cells [30,31]. Importantly, the central residue of FMC63 epitope is R163 and any mutation of the codon of R163 amino acid results in complete disruption of the epitope [30]. Additionally, the FMC63 epitope could also be disrupted when exon 2-encoded amino acid sequences are deleted [32]. Another factor that affects the *CD19* transcription and can also induce the appearance of truncated protein is intron 2 and intron 6 retention in mRNA as was revealed in lymphoma and B-ALL [29,33,34]. In fact, the shift towards higher intron 2 retention in therapy-resistant leukemias was apparent in almost 80% of paired B-ALL samples [35]. Later studies also showed that *CD19* splice variants lacking exon 2 were detected in B-ALL pediatric patients at diagnosis and could be the cause of initial leukemia resistance to CD19 CAR-T treatment [36]. Additionally, it was reported that preexisting CD19-negative subpopulation or progenitors detected in B-ALL patients before the CD19-CAR-T therapy could contribute to the failure and relapse after therapy administration [37,38]. These antigen-negative subclones become dominant within the tumor after immune selection pressure. A retrospective analysis of 628 B-ALL samples revealed that approximately 17 % of patients had CD19-negative tumor cells before treatments [39]. Moreover, some epigenetic mechanisms can affect the immunotherapy outcome. It was shown that in chronic lymphocytic leukemia (CLL) and Burkitt lymphoma antigen loss following CD19 CAR-T treatment was due to hypermethylation of the *CD19* promoter in tumor cells [40].

Many studies have shown distinct functional properties of CAR-T cells incorporating either 4-1BB or CD28 costimulatory domains [41,42]. 4-1BB CARs reveal longer persistence and higher proliferative ability [41,43], while CD28 CARs exhibit more rapid tumor elimination and higher potency for eradication of antigen-low tumors [44,45]. Moreover, 4-1BB CARs demonstrate a higher capability of trogocytosis-mediated antigen downregulation than CD28 CARs [46]. Furthermore, in the real-world clinical comparisons, CD28-based axi-cel demonstrated significantly improved responses and longer progression-free survival than 4-1BB-based tisa-cel in lymphoma patients, remaining a bit more toxic [47–50]. In the case of B-ALL, the differences between the efficacy of 4-1BB- and CD28-based products are not so evident, but tisa-cel remains the most often used, which can make the clinical data comparison more difficult [51,52]. Although extensive research about these products has been conducted, the knowledge of how the different clinical constructs can shape the process of antigen-dependent resistance development is still poorly understood. Therefore, in this study, we directly compared 4-1BB CARs and CD28 CARs to investigate the role of the costimulatory domain in CD19 CAR-T therapy resistance development in two distinct B-cell malignancy models.

## Results

### 4-1BB- and CD28-based CD19-CAR constructs differentially affect the dynamic of CD19 antigen downregulation and sensitivity to CD19 CAR-T cells

To understand the role of the costimulatory domains in the development of resistance, we generated *in vitro* models of B-cell lymphoma (Raji) or B-ALL (Nalm-6) cells exposed to either 4-1BB or CD28 FMC63-based CD19 CAR-T cells (Fig.1A, left panel). To mimic the clinical situation as closely as possible, we used the CAR structures identical to the structure of Kymriah (tisa-cel, CD19-4-1BB) and Yescarta (axi-cel, CD19-CD28) CAR clinical products (Fig. 1A, right panel; [1,2]). To establish the models, tumor cells were co-cultured side by side for 24 h with unmodified T cells (targets further referred to as control (con)), CD19-4-1BB CAR-T cells (targets referred to as CAR-4-1BB (4-1BB)) or CD19-CD28 CAR-T cells (targets referred to as CAR-CD28 (CD28)) for 19 subsequent expositions (passages). For every passage, the same conditions were used with modified T cells expressed comparable CAR levels (Fig. 1B). Raji cells were co-cultured with effectors in E:T 0.25:1 or 0.5:1, and Nalm-6 in E:T 0.1:1 or 0.25:1, depending on the passage number. The E:T ratio were experimentally selected to achieve the suboptimal effectors’ dose which resulted in elimination of approximately 70-80% targets at the initial stages. After each contact of tumor cells with CAR-T cells, tumor cells were harvested and monitored for CD19 levels with HIB19 clone antibody using flow cytometry (Fig. 1C). These experiments revealed a distinct pattern of changes in CD19 surface expression in tumor cells exposed to either CD19-4-1BB or CD19-CD28 CAR-T cells. While CD19-4-1BB CAR-T cells led to profound surface CD19 downregulation or almost its complete loss in Nalm-6 and Raji cells, respectively, exposure to CD19-CD28 CAR-T cells only partially downregulated CD19 surface levels (Fig.1C).

**Figure 1.**
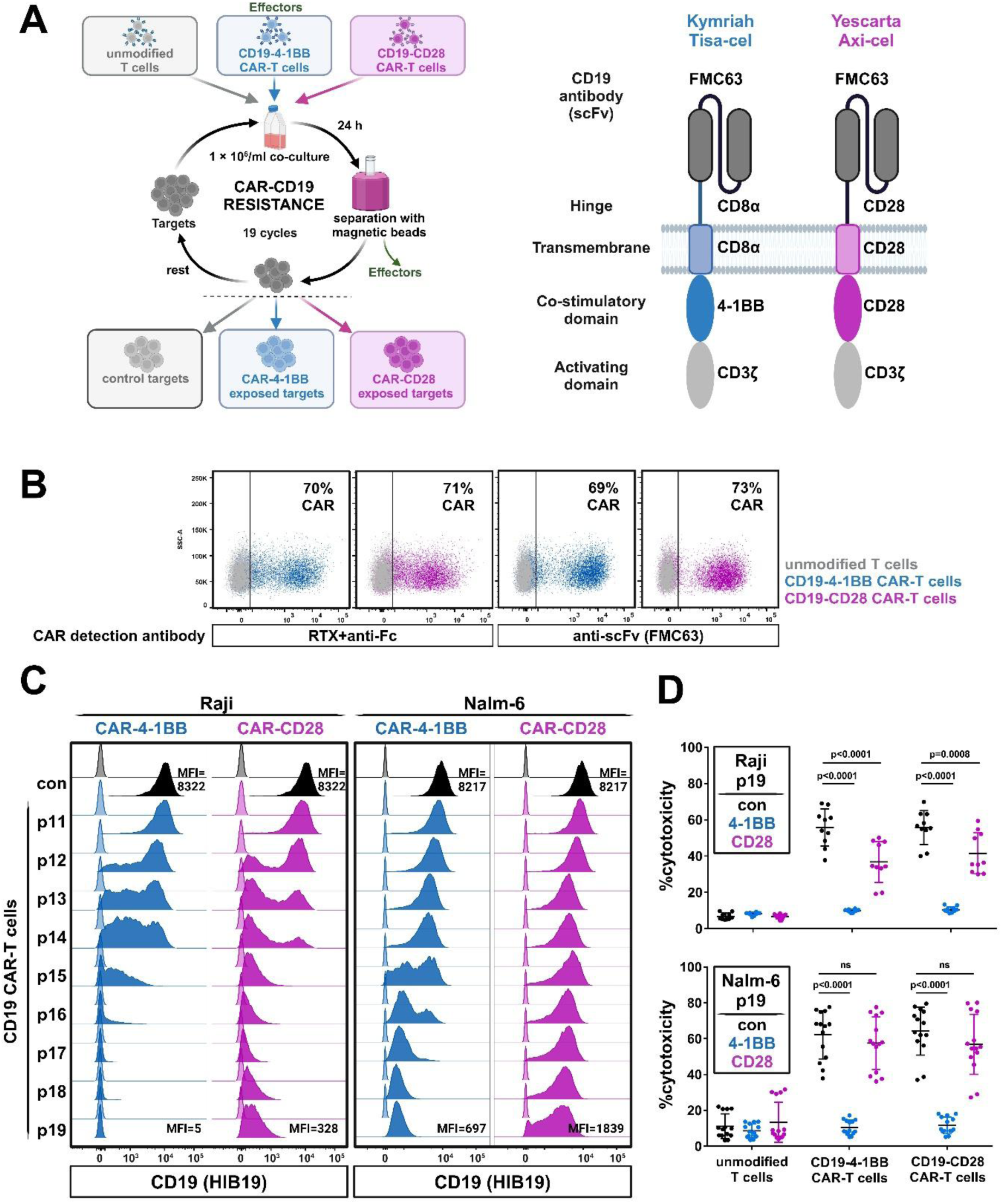
CAR-CD19 costimulatory domain differently affects the process of CD19 loss and sensitivity to effectors in lymphoma and B-ALL *in vitro* models of CD19 CAR-T resistance. (**A**) Scheme of procedure used for generation of resistant models (left panel). Lymphoma (Raji) and B-ALL (Nalm-6) models were established through 19 subsequent and parallel expositions to effectors (unmodified T-cells, CD19-4-1BB- or CD19-CD28 CAR-T cells), in the same conditions and E:T ratio for every passage. The structures of CAR-CD19 constructs bearing 4-1BB or CD28 costimulatory domain (right panel) were identical to clinically used product tisa-cel and axi-cel, respectively. (**B**) Efficacy of T-cells transduction with CAR-CD19-4-1BB and CAR-CD19-CD28 constructs. Detection of CAR-positive T-cells (a representative plot for one of the PBMCs’ donors) was performed by flow cytometry staining with RTX followed by secondary AlexaFluor 647-conjugated anti-Fc anibody (left) or anti-scFv (FMC63) antibody (right). (**C**) Flow cytometry analysis of CD19 surface expression in Raji cells (left panel) and Nalm-6 cells (right panel) during the process of long-term co-culture with CD19 CAR-T cells bearing either 4-1BB or CD28 costimulatory domain. The CD19 expression was detected by the HIB19 antibody clone after targets-effectors co-culture separation and targets recovery (up to 10 days). Raji or Nalm-6 cells co-cultured with unmodified T-cells are shown in black (con), co-cultured with CD19-4-1BB CAR-T cells in blue (Raji/Nalm-6 CAR-4-1BB) and with CD19-CD28 CAR-T cells in magenta (Raji/Nalm-6 CAR-CD28). The staining of anti-CD19 antibody (filled histograms) was compared to isotype controls (transparent histograms). p11-p19 reffers to the passage number. (**D**) Flow cytometry-based analysis of sensitivity of Raji (upper panel) and Nalm-6 (lower panel) cells to unmodified T-cells, CD19-4-1BB CAR-T cells or CD19-CD28 CAR-T cells. In the experiment, control (con) Raji and Nalm-6 target cells and their counterparts after long-term co-culture (19 passages) with CD19 CAR-T cells incorporating 4-1BB or CD28 domain were used. The plot shows the % cytotoxicity of effectors against targets. Data are presented as the mean ± SD from 5-7 donors analyzed in 2 technical repeats each. Statistical analysis was performed using two-way ANOVA with interaction and Tukey’s post-hoc test for multiple comparisons. Only selected significances were shown on the plots.

We then analyzed how CD19 downregulation affects tumor cells’ sensitivity to CAR-T cells. Consistent with CD19 loss, Raji cells exposed to CD19-4-1BB CAR-T cells (Raji CAR-4-1BB) were completely resistant to CD19 CAR-T cells incorporating either 4-1BB or CD28 costimulatory domains (Fig.1D, upper panel). Surprisingly, Nalm-6 exposed to CD19-4-1BB CAR-T cells (Nalm-6 CAR-4-1BB), although slightly positive for CD19, as detected by the HIB19 clone (Fig. 1C), also displayed full resistance to CD19 CAR-T cells (both 4-1BB and CD28 variant) (Fig.1D, lower panel). Conversely, CD19-CD28 CAR-T cells did not induce a full resistance of tumor cells. While CAR-CD28-exposed Raji cells demonstrated a significantly decreased susceptibility to either 4-1BB-based or CD28-based CD19 CAR-T cells, the sensitivity of CAR-CD28-exposed Nalm-6 cells was barely affected (Fig. 1D).

From these data, we concluded that CD19 CAR-T cells bearing either 4-1BB or CD28 costimulatory domain affect the dynamic of CD19 antigen loss differently, with various consequences for resistance development in both lymphoma and leukemia cellular models.

### Long-term co-culture with CD19 CAR-T effectors reveals different patterns of CD19 loss in lymphoma and B-ALL

To understand this phenomenon, we performed surface CD19 staining with both HIB19 and FMC63 clone in Raji and Nalm-6 models after 19 exposures to CD19-4-1BB and CD19-CD28 CAR-T variants. We detected the complete loss of FMC63 epitope in both Raji and Nalm-6 exposed to CD19-4-1BB CAR-T cells (Fig. 2A), explaining their full resistance to the therapy.

**Figure 2.**
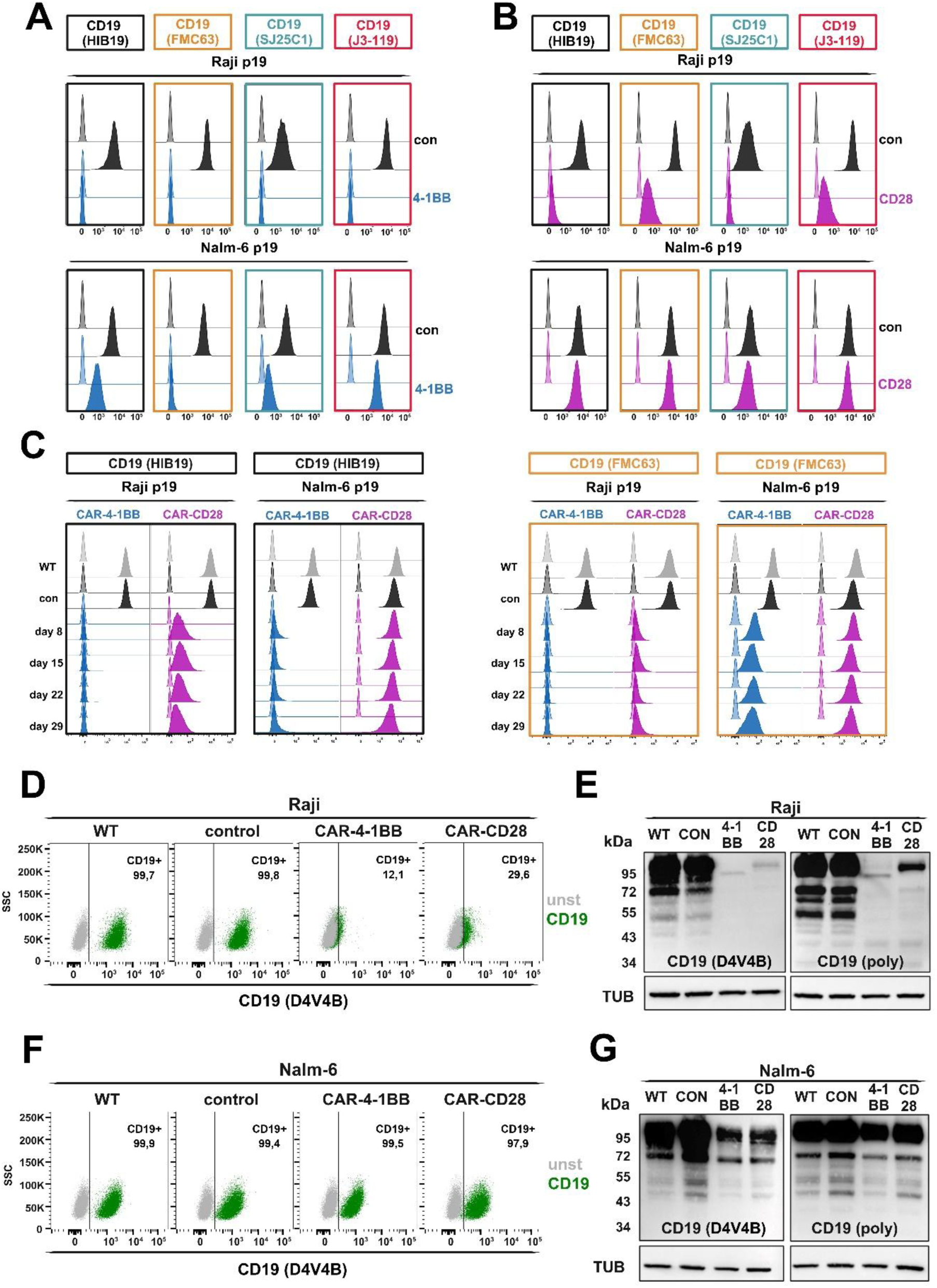
CD19-4-1BB CAR-T cells, but not CD19-CD28 CAR-T cells induce complete resistance in Raji and Nalm-6 models with different mechanisms of CD19 loss. (**A-B)** Flow cytometry analysis of CD19 surface level with different anti-CD19 antibodies clones – HIB19 (black), FMC63 (orange), SJ25C1 (turquoise) and J3-119 (red) in Raji and Nalm-6 control (con, black) and after long-term co-culture (19 passages) with effectors - CD19-4-1BB CAR-T (**A**, blue) or CD19-CD28 CAR-T cells (**B**, magenta). The staining of anti-CD19 antibody (filled histograms) was compared to isotype controls (transparent histograms). (**C)** The stability of resistance phenotype on Raji and Nalm-6 CAR-4-1BB and CAR-CD28 models was evaluated by flow cytometry CD19 surface level analysis with HIB19 clone (left panel) and FMC63 clone (right panel) after a culture of targets without effectors (8, 15, 22 and 29 days starting from day 0 – establishment of the models. (**D, F)** Flow cytometry-based analysis of CD19 intracellular epitope (clone D4V4B recognizing residues surrounding Leu427) level in Raji (**D**) and Nalm-6 (**F**) WT, control, CAR-4-1BB and CAR-CD28 tumor cells. Data show the representative experiment from 2 biological replicates. (**E, G)** Western blotting analysis of total CD19 protein level in Raji (**E**) and Nalm-6 (**G**) WT, control, CAR-4-1BB, and CAR-CD28 tumor cells evaluated by two antibodies recognizing different CD19 intracellular epitopes – residues surrounding Leu427 (clone D4V4B) or C-terminus residues (polyclonal antibody). Tubulin (TUB) was used as a loading control.

Differences in CD19 level detected by FMC63 and HIB19 clones prompted us to evaluate other commercially available anti-CD19 antibodies’ clones – SJ25C1 and J3-119 (Fig.2A, Suppl. Fig.1A). While in Raji CAR-4-1BB the profound loss of all evaluated epitopes was detected, in Nalm-6 CAR-4-1BB only the FMC63 epitope disappeared and three other CD19 epitopes were preserved at relatively high levels (Fig.2A, Suppl. Fig. 1A). A similar analysis upon CD19-CD28 CAR-T cells treatment led to different results. FMC63 epitope was detected at decreased, but still relatively high, levels in Raji CAR-CD28 (Fig.2B, upper panel) and almost totally preserved in Nalm-6 CAR-CD28 cells (Fig.2B, lower panel). These observations were in full accordance with the preserved sensitivity of these cell lines to CAR-T cells (Fig. 1D). Other tested epitopes, as detected with SJ25C1 and J3-119 clones, were downregulated to a similar extent as with HIB19 and FMC63 in Raji CAR-CD28 and only marginally changed in Nalm-6 CAR-CD28 (Fig. 2B, Suppl. Fig. 1A). Importantly, the phenotype of generated *in vitro* models was stable, as no fluctuations of CD19 levels were observed either with HIB19 (Fig.2C, left panel) or FMC63 (Fig. 2C, right panel) antibody over prolonged (up to 29 days) culture without effectors.

To further understand the differences between generated models, we evaluated the CD19 levels with antibody against the intracellular CD19 domain (D4V4B clone) using flow cytometry and Western blotting (D4V4B clone and a polyclonal antibody). In Raji cells, the profound loss of intracellular CD19 epitope was observed in both 4-1BB and CD28 models, as compared with controls (Fig. 2D). Consistently, the evaluation of total CD19 levels by two different antibodies revealed radically reduced CD19 levels with shifts in CD19 isoforms (Fig. 2E). Intriguingly, the isoforms preserved in Raji CAR-CD28 corresponded to the full-length CD19 protein, whereas in Raji CAR-4-1BB the major isoform had a marginally lower molecular weight. In contrast, Nalm-6 models did not display alterations in intracellular CD19 epitope levels (Fig. 1F). The total CD19 levels in Nalm-6 CAR-4-1BB and Nalm-6 CAR-CD28 were slightly reduced in comparison to control cells, but without observable shifts in CD19 isoforms patterns (Fig. 1G).

From these experiments, we concluded that while 4-1BB-CD19 CAR-T promoted the resistance in both Raji and Nalm-6 cells, CD28-CD19 CAR-T cells did not induce the antigen escape. In Raji model the resistance to 4-1BB-CD19 CAR-T cells led to complete loss of CD19 protein, while in the Nalm-6 model it resulted in FMC63-specific epitope loss affecting the recognition by CD19 CAR-T cells.

### The resistance induced by CD19-4-1BB, but not CD19-CD28, CAR-T cells is caused by genetic aberrations in *CD19* gene

To identify mechanisms underlying the observed differences between CD19-4-1BB and CD19-CD28 CAR-T cells, we first performed RT-qPCR experiments. For assessment of *CD19* mRNA transcripts levels, we used 3 different *CD19*-specific pairs of primers covering the exon 1-2, 3-4 and 4-5 junctions (Fig. 3A). Our results revealed significant changes in *CD19* mRNA levels in both Raji CAR-4-1BB and Raji CAR-CD28 models, as determined with all three *CD19* primer pairs. Surprisingly, the *CD19* mRNA downregulation was even more pronounced in Raji CAR-CD28 than in Raji CAR-4-1BB (Fig. 3B). In contrast, Nalm-6 cells showed less pronounced changes in *CD19* mRNA levels in both CAR-4-1BB and CAR-CD28 model (Fig. 3C). Altogether, while transcriptional changes in both Raji and Nalm-6 models exposed to CD19-CD28 CAR-T cells fully corresponded to changes at the protein level, the mechanism underlying the resistance of models exposed to CD19-4-1BB CAR-T cells remained elusive. Therefore, we further performed deep whole-transcriptome bulk RNA sequencing (RNAseq) of all generated models. To investigate changes in the transcriptome between CAR-4-1BB and CAR-CD28 models in Raji and Nalm-6 cells, as well as to analyze the *CD19* transcript sequence, we performed differential gene expression analysis (DESeq2). This analysis confirmed our RT-qPCR results and revealed the significant and more pronounced downregulation of *CD19* transcript in CAR-CD28 (log_2_FC=-4.23) than in CAR-4-1BB (log_2_FC=-2.25) in Raji models (Fig. 3D, right panel). Simultaneously, *CD19* transcript levels remained unchanged in both Nalm-6 models (Fig. 3D). Moreover, *CD19* sequence analysis revealed point mutations in both Raji and Nalm-6 cells exposed to CD19-4-1BB CAR-T cells, but not in those exposed to CD19-CD28 CAR-T cells (Fig. 3E-F, Suppl. Fig. 2A-C). All observed mutations affected the FMC63 epitope coding region. Based on FMC63 epitope mapping performed by others [30,31], both exon 3 and 4 are indispensable for FMC63 epitope recognition (Fig. 3A). In our study, we observed in Raji CAR-4-1BB cells a single base deletion in exon 3 resulting in a frameshift (p.L156Cfs*25) just before the start of the FMC63 coding region, which caused premature termination of CD19 and thus elimination of the FMC63 epitope (Fig. 3E, Suppl. Fig. 2A). Furthermore, the variant allele frequency (VAF) for this mutation was 95% that together with the lack of CD19 at the protein level suggested the loss of heterozygosity (LOH) as a potential mechanism of silencing of the wild type allele. To investigate this, we analyzed the other single nucleotide polymorphisms (SNPs) across the *CD19* gene and in adjacent genes (NFATC2IP and SPNS1). We found one SNP in intron 2 of *CD19* (chr16-28932883-C-G, rs79018922) that was heterozygous in control and CAR-CD28 Raji cells, but homozygous (SNP not present) in the CAR-4-1BB variant (Suppl. Fig. 3A). The same pattern was observed in two neighboring SNPs in NFATC2IP (chr16-28951108-C-T, rs7201257) and SPNS1 (chr16-28982048-C-T, rs61747536) (Suppl. Fig. 3B-C) further confirming LOH at this locus. All these SNPs were categorized as benign/moderate benign [53–57]. Both identified events, frameshift mutation and LOH, fully agree with CD19-loss-dependent resistance of Raji CAR-4-1BB. Simultaneously, a distinct mutational background was observed in Nalm-6 CAR-4-1BB cells, in which two point mutations were identified: a frameshift (p.S103Lfs*27) with 30% VAF affecting exon 2 that was predicted to lead to a truncated CD19 protein in exon 3 (Fig. 3F, left panel) and a missense substitution (p.R163H) with 55% VAF affecting Arg163 located in exon 3 (Fig. 3F, right panel). It has been previously reported that amino acid residues W159, R163, K220, P222 located in exon 3 and exon 4 are crucial for FMC63 antibody binding with R163 being central and indispensable for this process (Fig. 3A, [30,31]). In the raw data reads from RNAseq both mutations were not mapped in the same read-pairs, thus remaining in the trans configuration (Suppl. Fig. 3D). Considering the complete loss of FMC63 epitope in Nalm-6 cells (Fig. 2A), two different clones harboring different mutations may have emerged in Nalm-6 upon exposition to CD19-4-1BB CAR-T cells. Finally, in all Raji and Nalm-6 variants, we consistently detected a SNP (p.L174V) with 100% VAF (Suppl. Fig. 2D), which is categorized as a benign/strong benign polymorphism (chr16-28933075-C-G, rs2904880) by a VarSome [53] database and variety of algorithms [54–57] and was also detected in lymphoma patients treated with 4-1BB- and CD28-based CAR-T cells [58,59].

**Figure 3.**
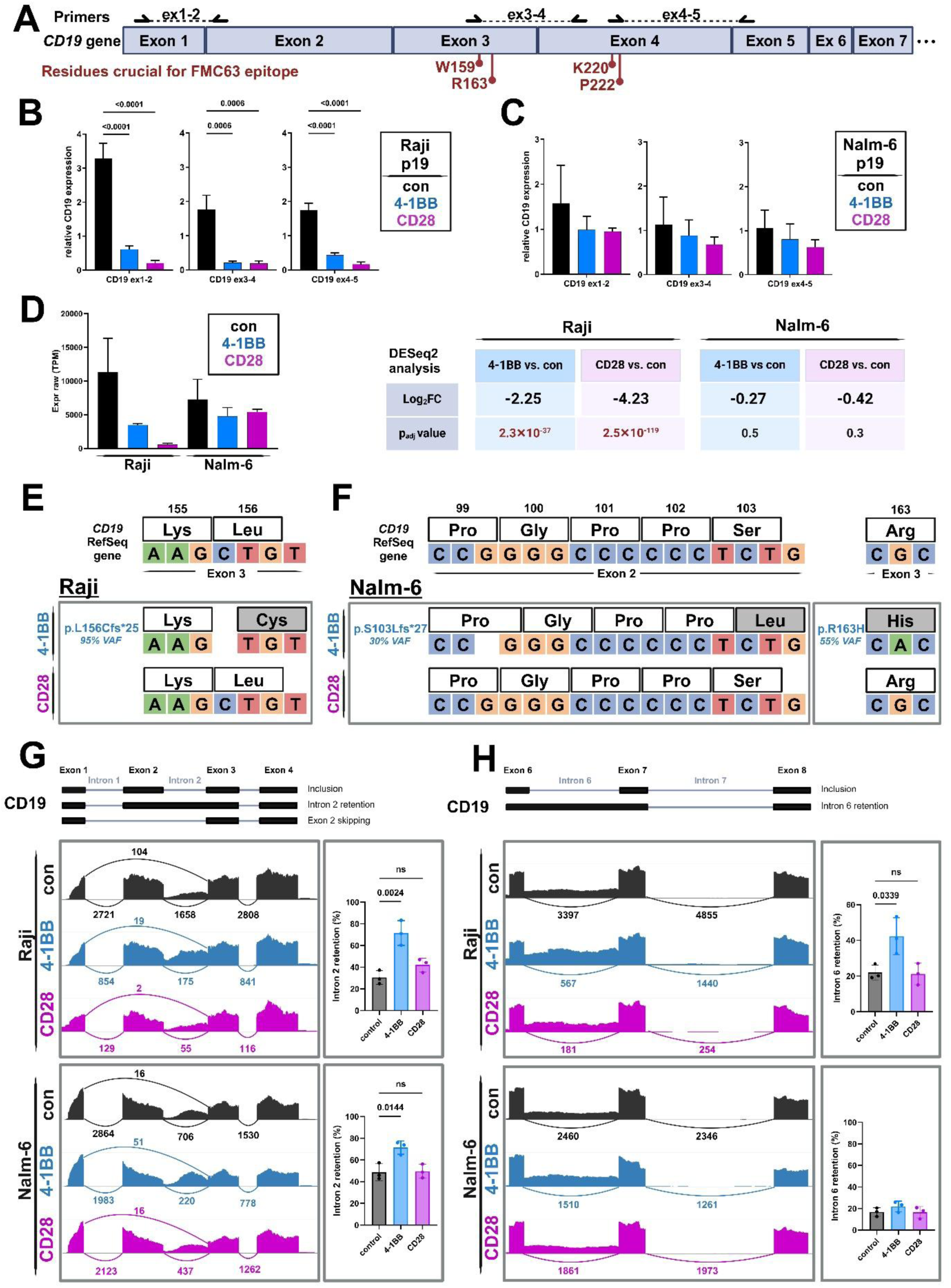
Repeated exposures of tumor cells to CD19-4-1BB CAR-T cells, but not CD19-CD28 CAR-T cells, result in genetic defects in *CD19* gene. (**A)** Scheme of *CD19* gene fragment (exon 1 to exon 7; grey blocks) together with the location of the primers used for RT-qPCR experiments (ex1-2, ex3-4, ex4-5, black arrows) and codons coding amino acid residues (W159, R163, P222, K220) essential for FMC63 epitope recognition. (**B-C)** RT-qPCR experiments showing relative *CD19* mRNA level in Raji (**B**) and Nalm-6 (**C**) control (con) cells and after 19 passages of effectors’ exposures to CD19-4-1BB-CAR-T (4-1BB, blue) or CD19-CD28-CAR-T-cells (CD28, magenta). The relative mRNA expression was assessed by 3 different pairs of *CD19*-specific primers covering exon 1-2, exon 3-4 or exon 4-5 junctions. The plot shows the CD19 mRNA level calculated with ΔCt method relative to the mean of TBP and GUSB as housekeeping genes. Data are presented as the mean ± SD from 3-4 biological repeats performed in technical triplicates. Statistical analysis was performed using one-way ANOVA with Tukey’s post-hoc test for multiple comparisons. (**D)** RNAseq analysis of CD19 transcript level in Raji and Nalm-6 control (con) and after exposition (19 passages) to CD19-4-1BB CAR-T cells (4-1BB) or CD19-CD28 CAR-T cells (CD28). The plot (left panel) shows raw expression (transcript per million, TPM) of CD19 transcript. Data is presented as mean ± SD from 3 biological replicates. Right panel shows the results of differential gene expression analysis (DESeq2) – the log_2_FC (fold change) and the p-value between control and 4-1BB or CD28 variants in Raji and Nalm-6 cell lines. (**E-F)** *CD19* sequence in Raji (**E**) and Nalm-6 (**F**) after exposition (19 passages) to CD19-4-1BB CAR-T cells (4-1BB) or CD19-CD28 CAR-T cells (CD28), as detected in RNAseq data and compared with CD19 RefSeq. The scheme shows the fragments of nucleotide sequences (blue – C, cytosine; orange – G, guanine; red – T, tymine; green – A, adenine) together with protein sequences, amino acid positions and exon numbers across the reference *CD19* gene and those detected in Raji and Nalm-6 CAR-4-1BB/CD28 variants. The results of mutations detected in 4-1BB cell lines (amino acids and together with variant allele frequency, VAF [%]) are marked in blue. (**G-H)** Intron 2 (**G**) or intron 6 (**H**) retention analysis in Raji and Nalm-6 control cells and after exposition (19 passages) to CD19-4-1BB CAR-T cells (4-1BB) or CD19-CD28 CAR-T cells (CD28) based on RNAseq data. Left panels show sashimi plots with coverage and the number of junction reads in Raji (upper panel) and Nalm-6 (lower panel) from pooled data of 3 biological replicates of each variant. The plots (right panels) show the frequency (% of all isoforms) of intron 2 (**F**) or intron 6 (**G**) retention calculated using the R/Bioconductor package ASpli. The data are presented as mean ± SD from 3 biological replicates. Statistical analysis was performed using one-way ANOVA with Tukey’s post-hoc test for multiple comparisons. Only selected significances were shown on the plots.

Moreover, after repeated exposures to CD19-4-1BB CAR-T cells, but not to CD19-CD28 CAR-T cells, we observed a significantly increased frequency of *CD19* isoform with intron 2 retention in Raji and Nalm-6 (Fig. 3G) and intron 6 retention in Raji cells (Fig. 3H). Both introns introduce the STOP codon and should result in truncated protein. Given that short-read sequencing is not well-suited for intron retention quantitation [60], we PCR-amplified the entire CD19 cDNA from Raji samples and subjected the PCR products to Oxford Nanopore Technologies long-read (lr) sequencing, as described by us previously [61]. Using this targeted lrRNA-seq approach, we were able to verify increased retention of both intron 2 and intron 6 (Suppl. Fig. 4). Additionally, we observed the tendency of exon 2 skipping in *CD19* junction’s events in Nalm-6 CAR-4-1BB (Fig. 3G, lower panel, 51 junction reads in 4-1BB vs. 16 in con/CD28). Collectively, prolonged exposure of either lymphoma or B-ALL cells to CD19-4-1BB CAR-T cells led to resistance dependent on genetic aberrations of *CD19* gene while CD19-CD28 CAR-T cells did not modify tumor cells’ mutational landscape.

Furthermore, to strengthen our observations, we compared our results with already available clinical data. Importantly, the same amino acid mutations (except for the one in codon L156), as well as CD19 intron retention isoforms detected in our 4-1BB models, were also described in r/r B-ALL/lymphoma patients treated with CD19 CAR-T cells (Table 1). Notably, in all reported cases patients were treated with 4-1BB-based CAR-T products, further validating our findings on the mechanisms of CD19-CAR-4-1BB-driven resistance.

**Table 1.** Comparison of mutation/intron retention (IR) detected in Raji and Nalm-6 *in vitro* models with available clinical data.

| CD19 mutation or IR in CAR-RES models | CD19 mutation/IR in r/r B-ALL/lymphoma patients |  |  | References |
| --- | --- | --- | --- | --- |
|  | Mutation | Disease | CAR-T therapy |  |
| **p.L156Cfs\*25** | Not detected | - | - | - |
| **p.S103Lfs\*27** | S103Lfs*27 | B-ALL | CTL019 (Tisa-cel) | [62] |
| **p.R163H** | R163L | B-ALL | CTL019 (Tisa-cel) | [29] |
|  | R163L | Lymphoma | BinD19 (CD19 CAR TCR-zeta/4-1BB) | [58] |
| **Intron 2 retention** |  | B-ALL | CTL019 (Tisa-cel) | [29,33,63] |
|  |  | B-ALL | N/A | [38] |
|  |  | Lymphoma | N/A | [34] |
|  |  | Lymphoma | Axi-cel | [64] |
| **Intron 6 retention** |  | Lymphoma | N/A | [34] |

### The phenotypic differences in tumor cells resistant to 4-1BB- and CD28-based CAR-T therapies are not limited to those targeting the CD19 antigen

To further investigate the differences between CAR-4-1BB and CAR-CD28 Raji and Nalm-6 models, we analyzed the RNA-seq data in the context of the changes across the whole transcriptome. We found 311 genes significantly upregulated and 480 significantly downregulated in Raji CAR-CD28 cells, while only 120 upregulated and 130 downregulated in Raji CAR-4-1BB in comparison to Raji control cells (Fig. 4A, upper panel). The commonly changed genes accounted for about 50% of all changes in Raji CAR-4-1BB and no more than 20% in Raji CAR-CD28. (Fig. 4A). On the other hand, in Nalm-6 cells, we found more significantly changed genes in CAR-4-1BB (522 upregulated, 694 downregulated) than in CAR-CD28 (298 upregulated, 417 downregulated), and only marginal number of these genes were commonly regulated (Fig. 4B). The differences between CAR-4-1BB and CAR-CD28 tumor cells variants were further confirmed by reactome signaling pathway analysis, both up- and downregulated (Suppl. Fig.5A-D, Suppl. Fig. 6A-D).

**Figure 4.**
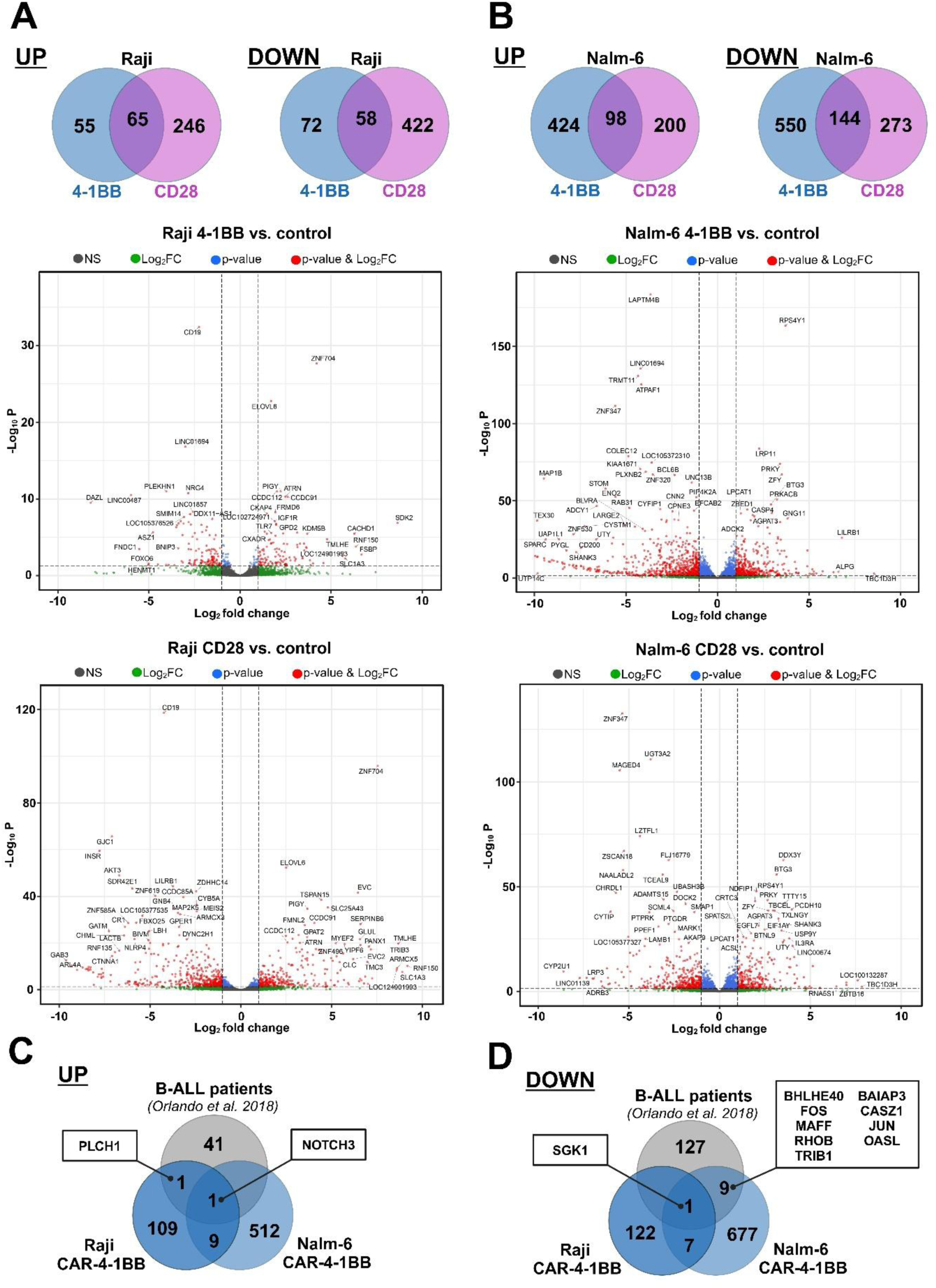
(**A-B**) Venn diagrams showing the comparison of significantly upregulated or downregulated genes between Raji CAR-4-1BB and CAR-CD28 (**A, upper panel**), and Nalm-6 CAR-4-1BB and CAR-CD28 (**B, upper panel**). The significantly changed genes (p-value > 0.05, log_2_FC > 1 (up) or log_2_FC < −1 (down)) were extracted based on differential gene expression analysis (DESeq2). Volcano plots showing the results of differential gene expression analysis (DESeq2) between Raji control and CAR-4-1BB (**A, middle panel**), Raji control and CAR-CD28 (**A, lower panel**), Nalm-6 control and CAR-4-1BB (**B, middle panel**) and Nalm-6 control and CAR-CD28 (**B, lower panel**). The cut-off point for p-value was set as 0.05 and for log_2_FC as 1 or −1. (**C-D)** Venn diagrams showing the comparison of significantly upregulated (**C**) or downregulated (**D**) genes between B-ALL patients treated with tisa-cel (re-analysis of RNAseq data from Orlando et al. (2018) and, Raji and Nalm-6 after long-term co-culture with CD19-4-1BB CAR-T cells (CAR-4-1BB). The significantly changed genes (p-value > 0.05; log_2_FC > 1 (up) or log_2_FC < −1 (down)) were extracted based on differential gene expression analysis (DESeq2) between relapsed samples vs. screening samples from Orlando’s data set and CAR-4-1BB vs. control samples in Raji and Nalm-6 models.

**Figure 5.**
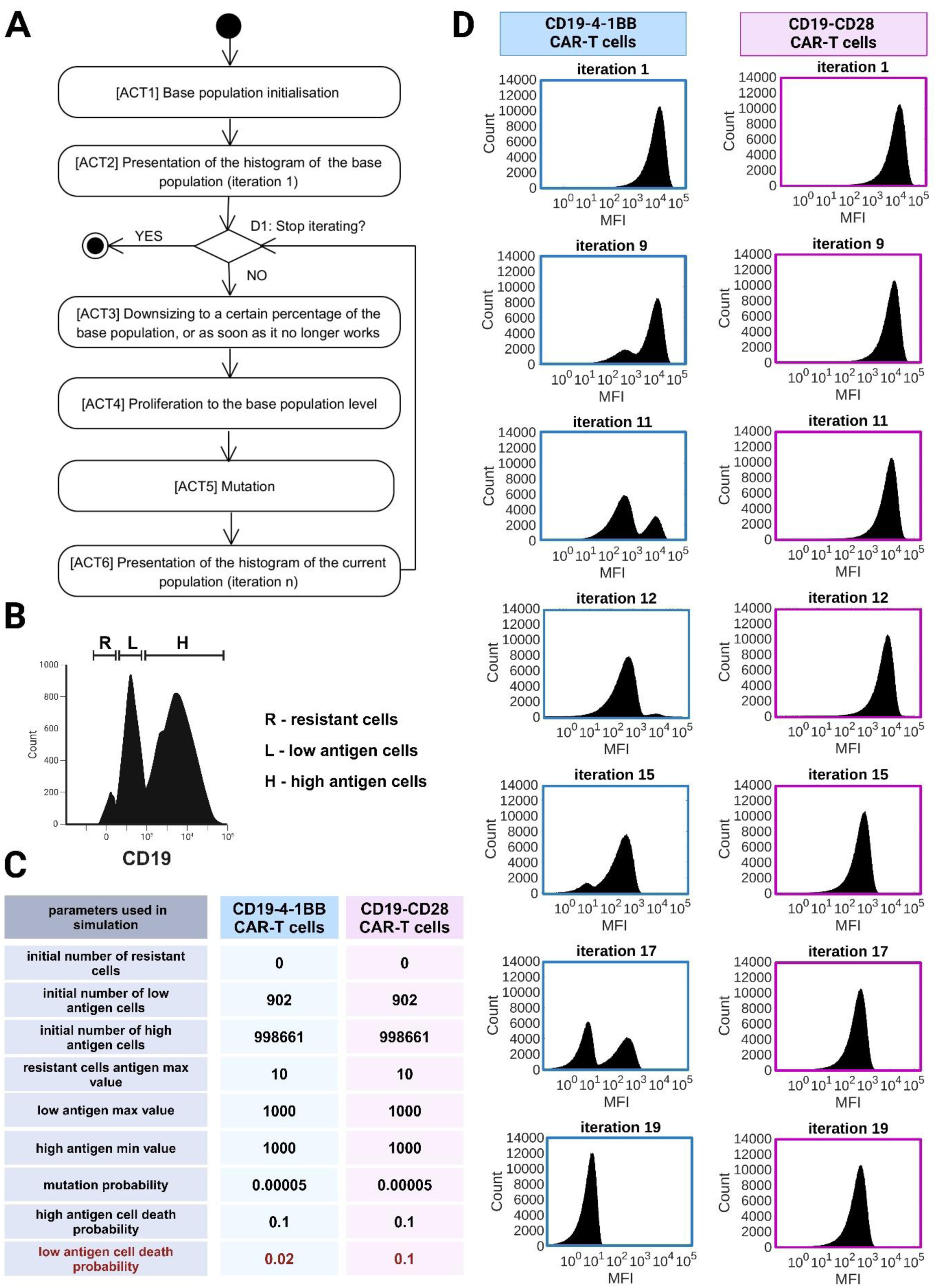
Mathematical simulation models of CD19 changes in tumor cells upon exposition to CD19-4-1BB CAR-T cells or CD19-CD28 CAR-T cells. (**A**) General simulation algorithm presented on activity diagram with the six subsequent experimentation stages. (**B)** Graphical representation on the histogram of the distinct subpopulations (resistant, low-antigen and high-antigen cells with defined antigen levels. (**C)** Main parameters set for simulations. (**D)** Histograms showing changes in CD19 MFI in subsequent passages (iterations) mimicking contact with CD19-4-1BB CAR-T cells (left) or CD19-CD28 CAR-T cells (right). The graphs show selected iteration from 19 that were performed.

Moreover, to further validate the transcriptomic changes observed in Raji and Nalm-6 models with clinical data, we re-analyzed the transcriptomic dataset from Orlando et al. 2018 [29]. In this dataset, 12 paired B-ALL patients’ samples were analyzed by RNA-seq at the stage of screening and after relapse. Since patients were treated with 4-1BB-based CD19 CAR-T therapy (tisa-cel), we compared this data with our 4-1BB-based resistant models only. Surprisingly, we detected only 189 significantly changed genes in patients’ data (43 upregulated and 137 downregulated) (Fig. 4C-D). In consequence, we identified only *NOTCH3* and *SGK1* as commonly changed across 3 data sets: patients’ samples, Nalm-6 CAR-4-1BB and Raji CAR-4-1BB. Notch3/4-signaling pathways, that were significantly enriched in Raji and Nalm-6 resistant models (Suppl. Fig.5-6 A-B) were previously reported to negatively correlate with B-ALL cells apoptosis [65].

In summary, repeated exposure to CD19-4-1BB or CD19-CD28 CAR-T cells induces phenotypic and transcriptomic changes beyond CD19 antigen loss, with 4-1BB-based models reflecting patterns seen in relapsed B-ALL patients.

### A mathematical simulation of resistance highlights the differences in how tumor cells are shaped by 4-1BB- or CD28-based CD19 CAR-T therapy

To explain the differential potential of 4-1BB-based and CD28-based CD19 CAR-T cells to induce resistance, we performed mathematical simulations (Fig. 5A). A base population of tumor cells, initially devoid of resistant cells, was defined and subjected to modifications using parameters that mimic the effects of 4-1BB- or CD28-based CD19 CAR-T cell therapy (Fig. 5B). The starting population was identical for both therapies and consisted of three subpopulations based on the antigen levels: resistant cells (CD19 MFI < 10), low-antigen cells (CD19 MFI between 10 and 1000) and high-antigen cells (CD19 MFI >1000) (Fig. 5B). In the simulations, the mutation probability for transition from high-antigen to low-antigen cells and from low-antigen to resistant cells was set at 0.00005, reflecting somatic mutation rates in tumor cells (Fig. 5C). The only parameter distinguishing the two therapies was the probability of killing low-antigen cells. While CD19-CD28 CAR-T cells were equally effective against both high- and low-antigen cells, the killing efficiency of low-antigen cells by CD19-4-1BB CAR-T cells was set five times lower (Fig. 5C). The simulations spanned 19 iterations, corresponding to 19 *in vitro* passages (Fig. 5D). CD19-4-1BB CAR-T cells led to an expansion of low-antigen cells starting at iteration 11 due to their reduced killing efficacy against this subpopulation. This expansion increased the likelihood of mutations generating resistant cells, which became detectable by iteration 15 and eventually dominated the population by iteration 19 (Fig. 5D, left panel). In contrast, CD19-CD28 CAR-T cells maintained a stable proportion between low- and high-antigen cells across iterations. Consequently, the number of low-antigen cells remained insufficient to produce resistant cells within the 19 iterations (Fig. 5D, right panel). The observed reduction in CD19 MFI in tumor cells treated with CD19-CD28 CAR-T cells was attributed to downregulation of CD19 antigen expression in high-antigen cells. This phenomenon, likely driven by epigenetic or other regulatory mechanisms, was incorporated into the simulation to replicate *in vitro* findings. These results underscore the critical role of targeting low-antigen tumor cells in mitigating resistance and highlight differences in the selective pressures exerted by 4-1BB- and CD28-based CAR-T cells.

Based on the simulation results, we concluded that CD19-4-1BB CAR-T cells alter the genetic landscape of tumor cells by fostering the expansion of the low-antigen population, ultimately leading to the emergence of resistant cells. At this stage, we hypothesized that resistant clones were absent in the tumor cell population prior to contact with CAR-T cells and emerged during exposure to CD19-4-1BB CAR-T cell therapy. To validate this hypothesis, we performed a simulation introducing a single resistant cell into a population of approximately 10 million non-resistant cells (Suppl. Fig. 7A). Despite the extremely low initial frequency of resistant cells – undetectable under *in vitro* or *in vivo* conditions – they dominated the population over the course of several iterations, regardless of whether CD19-4-1BB or CD19-CD28 CAR-T cells were applied, as demonstrated by histograms from selected iterations (Suppl. Fig. 6B). From this, we inferred that the resistant cells observed following CD19-4-1BB CAR-T cell treatment in our *in vitro* models were not pre-existing in the untreated tumor population. If resistant clones had been present initially, both CD19-4-1BB and CD19-CD28 CAR-T cells would have resulted in the emergence of a resistant phenotype after 19 treatment passages.

In summary, combined findings from *in silico* and *in vitro* models suggest that CD19-4-1BB CAR-T cells, but not CD19-CD28 CAR-T cells, generate resistant cells through suboptimal elimination of low-antigen tumor cells. This process appears to depend on the introduction of *de novo* mutations in the *CD19* gene, which were absent prior to CAR-T cell exposure.

## Discussion

In the present study, we demonstrate that prolonged exposure to CD19-4-1BB CAR-T cells leads to resistance of tumor cells to CD19 CAR-T therapy, while CD19-CD28 CAR-T cells do not induce this effect. While it was already demonstrated by others that CD19-CD28 CAR-T cells outperform CD19-4-1BB CAR-T cells against antigen-low tumors [45], our study provides additional evidence of their superiority in the context of resistance. Accordingly, tumor cells exposed to CD19-4-1BB CAR-T cells, show *CD19* mutations (missense, frameshifts) and intron retentions not detected in the control cells before the treatment nor in cells exposed to CD19-CD28 CAR-T cells. The genetic changes in cells exposed to CD19-4-1BB CAR-T cells led to either selective FMC63 epitope or total CD19 protein loss in both created models of B-ALL (Nalm-6 cell line) and B-cell lymphoma (Raji cell line). FMC63 epitope is the most widely studied epitope within CD19 antigen [66,67]. It has been already demonstrated by others that three CD19 residues (R163, K220, P222) are the top most significant contributors to interaction with FMC63, but also with other antibodies such as SJ25C1. Our findings our with agreement on the role of R163 as a determinant for FMC63 binding, as demonstrated by structural studies [67] and deep mutational scanning [66]. However, we have not observed the disruption of SJ25C1 binding in cells with missense R163H mutation.

To better understand the observed differences in the development of resistance between tumor cells exposed to CD19-4-1BB or CD19-CD28 CAR-T cells we performed mathematical simulation experiments. We observed that differences in the probability of killing low CD19 antigen tumor cells produces a different tumor phenotype mimicking *in vitro* results, where antigen-low tumor cells cause antigen escape-mediated resistance upon exposition to CD19-4-1BB CAR-T cells. While it is not entirely clear from our study how this process is advancing over time, we hypothesize that resistance to CD19-4-1BB CAR-T cells develops following a Lamarckian adaptive evolution. In this scenario in general, environmental factors cause genomic and heritable changes (mutations) that are targeted to a specific gene and provide adaptation to the original causative factor. It is obvious that the adaptive reaction to a specific environmental factor must be mediated by a molecular mechanism that results in the genomic change. Since CAR-T cells are not considered a mutagenic therapy, it is very unlikely that CD19-4-1BB CAR-T cells induce specific *CD19* mutations directly. However, tumor cells and cell lines are widely reported to have the increased spontaneous mutation rates [68]. In this context, we hypothesize that somatic mutations in malignant B cells are occurring during the subsequent passages and expositions to CAR-T cells. While mutations of the *CD19* gene in low-CD19-antigen cells can inevitably produce resistant cells, the probability of generation of resistant cells by CD19-high population is very low. This scenario aligns well with Knudson’s two-hit hypothesis for tumorigenesis. According to this theory, in order for a particular cell to become cancerous, both alleles of the tumor suppressor gene must be inactivated. In the case of our CAR-T-resistant models, both alleles of *CD19* gene need to be dysfunctional for the tumor cell to become resistant. In this context, the development of resistance to therapy resembles the process of cancer development. We hypothesize that in low-antigen population already one copy of *CD19* is somehow defected/inactivated and upon contact with CD19-4-1BB CAR-T cells a heterozygous cell from expanded low-antigen population receives a second hit in its remaining functional copy of *CD19* gene, thereby becoming homozygous for the mutated gene. Accordingly, in Raji CAR-4-1BB model, we have identified both frameshift mutation and loss of heterozygosity. In Nalm-6 cells we observed frameshift and missense mutations, but the detailed evolutionary trajectory is not clear and requires further studies.

Importantly, we have not identified any of these mutations in the control population of tumor cells before contact with CAR-T cells. While we are aware of the limitations of the power of the detection of single cells, our RNA sequencing depth allowed for the detection of approximately 0.1% population. Therefore, we also performed mathematical simulations with the introduction of one resistant cell into the source population of tumor cells. We observed that even with as low frequency as one resistant cell in the initial population of 10 million cells, resistant cells would be inevitably and successfully selected by both variants of CAR-T cells after 19 passages, assuming similar proliferation rates of resistant and sensitive cells. On the assumption that Darwinian selection is the process contributing to drug resistance in our models, it would inevitably lead to resistance development to both CD19-4-1BB CAR-T cells and CD19-CD28 CAR-T cells. Therefore, we excluded the possibility of a scenario in which purely Darwinian mechanisms of evolution and selection of malignant B cells containing mutations already present before the treatment underlie the differences in resistance development between 4-1BB- and CD28-CAR-T cells.

Changes observed in our models can mirror diverse scenarios encountered in patients treated with CAR-T-cells, in whom the escape of the tumor cells from CAR-T cells immunosurveillance could be achieved even faster and be dependent on both selection and phenotypic plasticity of tumor cells under pressure from CAR-T therapies. This may result from higher tumor heterogeneity in patients and the presence of not only CD19-low cells but also of negative clones [38]. Furthermore, our results clearly show that subsequent contacts of CAR-T cells with tumor cells increase the probability of antigen-negative escape. It remains in accordance with clinical data, where maintained persistence of CD19-4-1BB CAR-T cells in patients and their sustained pressure is associated with complete loss of the surface epitope recognized by the CAR and is more frequently observed for 4-1BB- than for CD28-containing CARs [28,29]. Consistently, in the real-world clinical comparisons in lymphoma patients, CD28-based axi-cel demonstrated significantly improved responses and longer progression-free survival than 4-1BB-based tisa-cel [47–50].

The advantage of our models above other resistant models produced so far with CRISPR/Cas9 library [69–71] or with artificial expression of CD19 [45] is that we generated resistant cells upon CAR-T cells pressure that can reflect natural physiology and clinical experience. Although our models were generated *in vitro* with two cell lines exposed to CD19 CAR-T cells with restricted E:T ratio, they thoroughly reproduce clinical findings. Accordingly, they demonstrate some changes already linked with primary/acquired resistance to the CD19-CAR-T therapy in patients, including selective FMC63 loss, specific *CD19* gene mutations and an increased frequency of *CD19* aberrant versions with intron 2 and/or intron 6 retention, as shown in Table 1. In all examined Raji and Nalm-6 variants, we also detected a benign polymorphism (p.L174V), which does not seem to be the direct cause of resistance. However, its role in recognition of FMC63 epitope by CD19 CAR-T cells cannot be excluded and requires further studies as this polymorphism was also reported in patients after CD19 CAR-T therapy [58,59].

Being aware of some limitations of our study, we also believe that our observations and conclusions can contribute to better recognition of consequences for patients treated with CAR-T therapies. Firstly, it becomes clear that patients can develop resistance to CD19-4-1BB-CAR-T-cells during the treatment, with no preexisting resistant clones. This is also in accordance with clinical data published by Orlando *et al.* [29], where mutations were not detected in patients before the treatment, despite exhaustive and detailed genomic studies performed prior to therapy. Considering this, it becomes apparent that it is not possible for CD19-4-1BB-CAR-T-cells-treated patients to predict their response to treatment and chances of relapse. Secondly, although CD19-4-1BB CAR-T cells have advantages in the clinics over CD19-CD28 CAR-T cells due to their high persistence as generally reported as a positive feature [41,43], it can also negatively impact the patients’ outcome in the long-term perspective. In general, the prolonged exposition of tumor cells to CD19-4-1BB CAR-T cells can suboptimally shape the tumor population and reinforce the adaptations in tumor cells that can be readily propagated by genetic inheritance. Finally, our observations on selective FMC63 epitope loss underlie the need to improve diagnostic procedures of B-ALL patients treated with CD19-immunotherapies that are currently performed with various anti-CD19 clones. Although SJ25C1 antibody and the HIB19 monoclonal antibody are reported to recognize overlapping epitopes, they are not mapped in details. Moreover, FMC63 and SJ25C1 monoclonal antibodies target the same site on CD19 albeit with different binding orientations and affinities [67]. In light of this, only detection of the FMC63 epitope, as it was shown by some clinical cases [72], delivers reliable information on the levels of target antigen accessible for commercial FMC63-based CD19 CAR-T cells and this simple and low-cost procedure should be included in the flow cytometry diagnostics panels.

## Materials and methods

### Cell lines

The cell lines used in this study were purchased from the German Collection of Microorganisms and Cell Cultures (Nalm-6) and the American Type Culture Collection (Raji, HEK293T). The Raj and Nalm-6 cells were cultured in RPMI-1640 medium (Gibco™, ThermoFisher) supplemented with 10% fetal bovine serum (FBS) (HyClone™, Cytiva), and 1% antibiotics – penicillin/streptomycin (Gibco™, ThermoFisher). HEK293T cells (RRID: CVCL_0063) were cultured in DMEM medium (Sigma-Aldrich) supplemented with 10% FBS. All cultures were maintained in a humidified atmosphere containing 5% CO_2_. All cell lines were routinely tested for *Mycoplasma spp* and only cells free of contamination were used.

### CAR construction

The CAR-CD19 (FMC63) construct in retroviral SFG vector bearing the CD8 hinge and transmembrane domain, 4-1BB costimulatory domain, CD3ζ signaling domain, and rituximab recognized-RQR8 epitope followed by T2A sequence in frame with CAR sequence (CAR CD19-4-1BB) previously used in [73] was obtained as a kind gift from Martin Pule (UCL, UK). The CAR-CD19 construct incorporating CD28 hinge, transmembrane and costimulatory domain, and CD3ζ signaling domain was constructed using the Gateway cloning system in retroviral pMP71 plasmid (CAR CD19-CD28). To unify the CAR system both mentioned above CAR-CD19 constructs were sub-cloned into the BamHI/SbfI restriction site of the lentiviral transfer plasmid pSEW [74], thereby replacing the GFP gene. Additionally to CAR CD19-CD28 the RQR8 epitope was added to help with CAR detection.

### Peripheral blood mononuclear cells (PBMCs) isolation and T cells selection

Peripheral blood mononuclear cells (PBMCs) were isolated from the whole blood of healthy donors obtained from the Regional Blood Center in Warsaw, Poland, with the waiver of consent granted by the Bioethics Committee of the Medical University of Warsaw, Poland. PBMCs were isolated from buffy coats with gradient centrifugation using Lymphoprep (STEMCELL). Subsequently, PBMCs were seeded onto the 6-well plate at 2 × 10^6^ cells/ml density in 3 ml RPMI-1640 medium supplemented with 10% FBS and 1% penicillin and streptomycin solution (RPMI full) and treated for 3 days with 1 µg/ml anti-CD3 (Thermo Fisher, Cat# 16-0037-81, RRID: AB_468854) and 1 µg/ml anti-CD28 (Thermo Fisher, Cat# 16-0289-81, RRID: AB_468926) antibodies to stimulate T cell activation. Then, the cells were modified by lentiviral transduction.

### Lentiviral T cell modifications and culture

To produce CAR viral particles, HEK293T cells were seeded at 6 cm plates (1 × 10^6^ cells per plate) and transfected using FuGENE HD transfection reagent (Promega) and transfer plasmid coding CAR construct simultaneously with VSV-G envelope expressing plasmid pMD2.G (RRID: Addgene_12259) and lentiviral packaging plasmid psPAX2 (RRID: Addgene_12260). For transfection, the 2.25:1 ratio of FuGENE HD reagent to DNA was used. After 48 h, the lentivirus-containing supernatant was harvested, centrifugated (500 × *g* at room temperature) and used directly for T cell transduction or frozen at −80°C for another transduction. For T cells transduction, 0.5 × 10^6^ of stimulated PBMCs were transferred on a 24-well plate and 1 ml of viral supernatant (or 1 ml of culture medium for unmodified T cells variant) and 4 µg/ml of polybrene (Sigma-Aldrich) was added. The cells were spinoculated with the viral particles for 1 h (1250 × *g* at 32°C). After spinoculation, the PMBCs were cultured in a humidified atmosphere containing 5% CO_2_. The next day, the second round of spinoculation with the fresh batch of viral supernatant was performed. After 6-8 h, the viral supernatants were replaced with the fresh portion of RPMI full medium supplemented with 200 U/ml IL-2 (PeproTech) and Dynabeads Human T-Activator CD3/CD28 (Thermo Fisher Scientific) at a bead-to-cell ratio of 1:3. After 3-5 days from transduction the CAR expression on T cells’ surface was evaluated by flow cytometry. After 5-6 days from transduction, the Dynabeads Human T-Activator CD3/CD28 beads were removed by magnetic separation, and the culture medium was changed to RPMI full medium supplemented with 100 U/ml IL-2. In these conditions, T cells were cultured for up to 6 weeks.

### CAR surface staining

To evaluate the modification efficacy with CD19-4-1BB and CD19-CD28 CAR constructs, the CAR-T-cells and unmodified T-cells were washed 3 times with PBS buffer and stained using one of two flow cytometry-based methods using BD FACSCanto II or BD LSRFortessa X-20 cytometers. In the first method, T cells were incubated with 100 µg/ml rituximab (RTX) following secondary staining with goat anti-human IgG, Fcγ fragment specific antibody conjugated with Alexa Fluor 647 (Jackson ImmunoResearch Labs Cat# 109-606-098, RRID: AB_2337899, dilution 1:200) detecting an RQR8 epitope expressed by CAR-modified T-cells. The % CAR positive cells was estimated based on the control variant of CAR-T cells incubated only with a secondary antibody. The second method for CAR expression detection was staining with anti-FMC63 scFv antibody conjugated with Alexa Fluor 647 (Cytoart, Cat# 200102, RRID: AB_2857946, dilution: 1:100) that specifically binds to the scFv region of CAR-CD19. The % CAR positive cells were estimated based on background from unmodified T cells stained with the antibody.

### CAR-CD19 resistance models obtaining

We used Raji and Nalm-6 cell lines, to generate the lymphoma and B-ALL resistance models of long-term co-culture with effectors, respectively. In the case of both cell lines, tumor cells were subjected to the subsequent and parallel passages with effectors (unmodified T cells, CD19-4-1BB CAR-T cells or CD19-CD28 CAR-T cells), in the same conditions for every passage. The target cells were seeded in co-culture with effectors cells (co-culture density 1 × 10^6^/ml) for 24 h at E:T 0.25-0.5:1 for Raji cells and 0.1-0.25:1 for Nalm-6 cells. For single passage the total number of target cells varied from 6 × 10^6^ to 40 × 10^6^ (depending on target cells condition). After 24 h of incubation, the cells were magnetically separated using the commercially available Human CD3 Positive Selection Kit (STEMCELL, Cat# 17851). The effector cells were removed and harvested target cells were left for the rest until the full recovery, and the exposure was repeated with the fresh portion of effectors. To generate Raji model, 6 different effectors’ donors were used, for Nalm-6 model – 5 different donors. Each donor was used for a maximum of 4 cycles. In summary, the cycle of targets-effectors 24 h co-culture was repeated 19 times for both Raji and Nalm-6 cell lines.

### Flow cytometry phenotyping

For flow cytometry phenotyping (CD19 antigen and other surface antigens), the 0.5 × 10^6^ of cells were washed twice in PBS Buffer, incubated with human Fc Block reagent (Becton Dickinson) for 15 min at room temperature, and stained with the appropriate amount of monoclonal antibodies (Suppl. Table 1) conjugated with fluorochrome for 20 min at room temperature. If needed, to assess viability, the cells were stained with Fixable Viability Stain 510 (Becton Dickinson) before the antibodies staining procedure. For intracellular staining of CD19 the membrane permeabilization was performed using Cytofix/Cytoperm kit (BD) before the staining with antibodies. Analyzes were performed using BD FACSCanto II or BD LSRFortessa X-20 cytometers.

### Cytotoxicity assays

For the CAR-T-cell-mediated cytotoxicity assay, the tumor target cells were stained with 0.5 µM CellTrace Violet kit (Thermo Fisher) and seeded with effector cells (unmodified T-cells or CAR-T-cells) at different effectors to targets ratio (E:T). After 18-24 h of incubation, the cells were stained with 0.5 µg/ml propidium iodide (PI) and immediately analyzed by flow cytometry to assess the cells’ viability. % cytotoxicity was calculated as % dead (PI-positive) cells from CTV-positive targets.

### Western blotting

Total protein lysates were prepared in the sample buffer (60 mM Tris-HCl pH 6.8, 2% SDS, 10% glycerol, 5% β-mercaptoethanol, bromophenol blue) followed by incubation at 95°C for 5 min. In each experiment, the same cells amount (3-4 × 10^6^) from all analyzed cell lines were used. Proteins from total cell lysates were separated in 8% (v/v) SDS-polyacrylamide gel. For electrophoresis always the same volume of prepared lysates was used (15 µl). Next, proteins were transferred onto nitrocellulose membranes and blocked with 5% (w/v) nonfat dry milk in TBST (Tris-buffered saline, pH 7.4 and 0.05% (v/v) Tween-20) for 1 h at room temperature. Then, the membrane was incubated overnight at 4°C with the following primary antibodies: anti-CD19 recognizing epitope surrounding Leu427 (Clone 1; 1:1000 dilution, Cell Signaling Technology, Cat #90176, RRID: AB_2800152), anti-CD19 recognizing C-terminus residues (Clone 2; 1:1000 dilution, Cell Signaling Technology, Cat #3574, RRID: AB_2275523) or anti-tubulin (1:1000, Sigma-Aldrich, Cat #CP06, RRID: AB_2617116). Tubulin was used as a loading control. Afterward, HRP-conjugated secondary antibodies were used for the detection of specific protein bands. Blots were exposed to the Super Signal chemiluminescent substrate (Thermo Fisher) and visualized using ChemiDoc Imaging System (Bio-Rad ChemiDoc MP Imaging System, RRID: SCR_019037).

### RT-qPCR

Total RNA from Raji/Nalm-6 cell lines samples was isolated using RNAsy Mini Kit (Qiagen) and RNase-Free DNAse Set (Qiagen) according to the manufacturer’s instructions. Next, cDNA synthesis was performed using 1 µg of RNA and Maxima First Strand cDNA Synthesis Kit for RT-qPCR (Thermo Fisher) and BioRad thermocycler. For qPCR reaction the PowerUp™ SYBR™ Green Master Mix (Applied Biosystems) was used together with three pairs of specific *CD19* primers (Suppl. Table 2). The reaction was run using a 7500 Fast Real-Time PCR System (Applied Biosystems). The relative *CD19* mRNA level was calculated using the comparative Ct (ΔCt) method relative to the mean of GUSB and TBP housekeeping genes.

### RNAseq and data analysis

Whole-transcriptome analysis was performed by deep bulk RNA sequencing (RNASeq). Each cell line variant was performed in three biological replicates. For cDNA library preparation 100 ng of total RNA was used in TruSeq Stranded Total RNA library preparation protocol with rRNA depletion (Illumina). Paired-end sequencing was performed (2×101 bp) with the required sequence depth of 30 millions pair-end reads (6 Gb) per sample. Sequencing was performed on the NovaSeq6000 instrument (Illumina).

Demultiplexing of the sequencing reads was performed with Illumina bcl2fastq (version 2.20). Adapters were trimmed with Skewer (version 0.2.2) [75]. The quality of the FASTQ files was analyzed with FastQC (version 0.11.5-cegat) [76]. Trimmed and cleaned reads were mapped onto the human genome (version GRCh38.110) to compute the counts in the form of Transcripts Per Million (TPM) using Salmon version 1.8.0 [77]. Prior to the calling of differentially expressed genes (DEG) similarity among samples were performed using a PCA analysis on regularized counts using the logarithm transformation as implemented in DESeq2 [78] to identify experimental covariates and batch effects among samples and replicates. Based on these analyses, the calling of DEGs was perform separately in each cell type, meaning Raji and Nalm-6 cells, comparing CAR-4-1BB and CAR-CD28 treated cells against control cells. The list of up- and downregulated DEGs were then identified using DESeq2 [78] at a p-value < 0.05 and a log_2_ Fold change above or below 1, respectively. Reactome [79] enrichment analyses based on significant DEGs was done using the enrichR R package version 3.2 [80].

The quantification of intro retention events on the CD19 gene in Raji and Nalm6 samples was done as follow. Reads were mapped onto the human genome (version GRCh38.110) using hisat2 version 2.2.1 [81] to obtain SAM files. SAM files were then converted to sorted and indexed BAM format using the SAMTOOLS version 1.18 [82]. Resulting BAM files containing the mapped reads were used to quantify the level of intron 2 and intron 6 retention using the ASpli R package version 2.12.0 [83].

### Oxford Nanopore Technologies (ONT) targeted re-sequencing

Total RNA from Raji cell lines was isolated using RNAsy Mini Kit (Qiagen) and RNase-Free DNAse Set (Qiagen) according to the manufacturer’s instructions. The cDNA was obtained from 1 µg of RNA using Maxima First Strand cDNA Synthesis Kit for RT-qPCR (Thermo Fisher) and BioRad thermocycler. Primers were designed to amplify the full-length coding region of CD19 by annealing to the 5’ (sense) and 3’ (antisense) exon present in all known isoforms. 10-ng aliquotes of cDNA were amplified using these primers with LongAmp Taq 2X Master Mix (New England Biolabs) for 29 cycles. The amplified cDNA was subjected to amplicon-seq (SQKNBD112.24, ONT) library preparation. The library was then loaded onto a Spot-ON flow cell R10.4.1 version (FLO-MIN114, ONT) and then sequenced in a MinION Mk1C device (ONT). Library was sequenced until there was at least 1000 reads per each sample multiplexed. After the library was sequenced, reads were aligned to the human genome (hg38) using Minimap2 version 2.24-r1122. Resultant BAM files were visualized in IGV version 2.19.1.

### *In silico* resistance models

To reproduce *in silico* the general phenomenon observed *in vitro* in cell lines exposed to either CD19-4-1BB CAR-T cells or CD19-CD28 CAR-T cells the MATLAB computing platform was used. The number of MATLAB scripts were written to: (i) prepare base population for its further evolution simulation, (ii) simulate the population evolution in subsequent passages (iterations), (iii) generate the histograms presenting population state for the particular iterations. This multi-stage approach supports the efficiency of experimentation by, among others, standardization of base population for various simulations and elimination of the need for re-simulation on the stage of histogram generation.

The activity diagram (Fig.5A) comprises experimentation stages:

[ACT1] - Base population is generated for the assumed number of cells with antigen value normal distribution with mean = 9000 and standard deviation = 3000. At this stage, the resistant cells maximum antigen value was set to 10. All of the resistant cells were removed from the base population during its generation (Fig. 5) or 1 cell per 10 million cells was introduced (Suppl. Fig. 6).

[ACT2] - The histogram for the base population presents the MFI basing on the cells antigen value. Measurement noise is represented by component generated with normal distribution with mean = 0 and standard deviation depending on the antigen value. For resistant cells standard deviation = 0, for other cells standard deviation = antigen value.

[ACT3] - The goal is to downsize the population to reach the reduction factor = 0.6. The reduction process is iterative, where in each iteration the particular cell can die with certain probability, depending on its antigen value. Death probability of high-antigen cells and low-antigen cells differ. The change in probability at the transition between low-antigen cells and high-antigen cells is linearized. Resistant cells are immortal, hence the resistant population is not downsized. In consequence, it is not possible to reach the reduction goal for populations with high number of resistant cells.

[ACT4] - The population is upsized to reach the count of base population. Each cell, regardless of antigen value, proliferates with the same probability.

[ACT5] - The high-antigen cells mutate to low-antigen cells and the low-antigen cells mutate to resistant cells with the same certain probability. The resistant cells do not mutate.

[ACT6] - In addition to [ACT2] the histograms generated for the subsequent iterations consider the reduced expression of antigen by introduction of sigmoid function that significantly reduces the MFI for high antigen cells in the neighborhood of iteration 12.

[D1] - The simulation process is stopped by the user.

### Statistical analysis and data visualization

The statistical analysis of obtained data was performed with the use of GraphPad Prism 7 and Statistica 13.3 software. The data was analyzed with parametric or non-parametric methods depending on data distribution. Assumptions of normality and homogeneity of variance, required for parametric methods, were verified using Shapiro-Wilk and Levene’s tests, respectively. In all analyses the tests were two-tailed and a p-value less than 0.05 was considered significant. Data were presented as mean value ± standard deviation (SD) or median with interquartile range (IQR) depending on data distribution. Figure legends provided detailed information about data presentation and analysis type. All plots were prepared using GraphPad Prism 7. Flow cytometry data were analyzed and visualized using FlowJo 10.9. Schemes and figures were created with BioRender.com.

## Acknowledgements

We would like to acknowledge Professor Wojciech Fendler, Dr. Akanksha Jaiswar and Mrs. Zuzanna Nowicka (Medical University of Lodz, Poland) for support in preliminary RNAseq data analysis and Else Marit Inderberg for cloning support. We thank Dr. Iwona Baranowska, Mrs Ewa Pięta and Mrs. Elżbieta Gutowska for routine PBMCs preparation and technical assistance. SW is grateful to the Flow Cytometry Core Facility at the Oslo University Hospital and Radiumhospitalets legater for their support.

