## Supplemental Figures for "The costimulatory domain influences CD19 CAR-T cell resistance development in B-cell malignancies"

**Suppl. Table 1**

| No. | Antigen | Clone | Fluorochrome | Catalog number | Manufacturer |
|---|---|---|---|---|---|
| **1** | CD19 | HIB19 | PECy7 | 25-0199-42 | Thermo Fisher |
| **2** | CD19 | HIB19 | BV421 | 562440 | BD |
| **3** | CD19 | HIB19 | PE | 12-0199-42 | Thermo Fisher |
| **4** | CD19 | FMC63 | PE | MAB1794H | Sigma Aldrich |
| **5** | CD19 | SJ25C1 | PE | 12−0198−42 | Thermo Fisher |
| **6** | CD19 | J3-119 | PE | A07769 | Beckman Coulter |
| **7** | CD19 | D4V4B | PE | 58960S | Cell Signaling Technology |

**Suppl. Table 2**

| Gene Name | Primer sequence (5'-3') | |
|---|---|---|
| **CD19 ex1-2** | Forward | GGAGAGTCTGACCACCATGC |
| | Reverse | ACTGCAGCACAGCGTTATCT |
| **CD19 ex3-4** | Forward | GAGCCCCAAGCTGTATGTGT |
| | Reverse | GGACACAGAGTCAGGGGGTA |
| **CD19 ex4-5** | Forward | AAGGGGCCTAAGTCATTGCT |
| | Reverse | CAGCAGCCAGTGCCATAGTA |
| **GUSB** | Forward | GAAAATATGTGGTTGGAGAGCTCATT |
| | Reverse | CGGAGTGAAGATCCCCTTTTTA |
| **TBP** | Forward | GCACAGGAGCCAAGAGTGA |
| | Reverse | GTTGGTGGGTGAGCACAAG |

**Supplementary Figure 1**

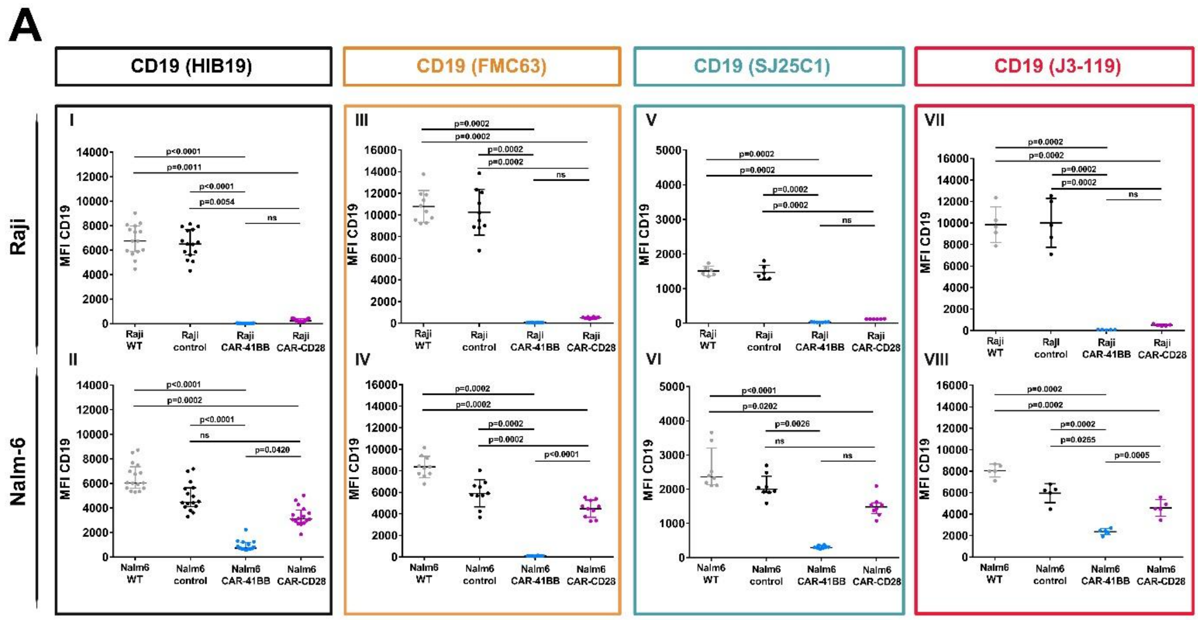


**Suppl. Figure 1**

(**A**) Flow cytometry-based analysis of CD19 surface level in Raji and Nalm-6 WT, control, CAR-4-1BB and CAR-CD28 cells assessed by different clones of anti-CD19 antibodies – FMC63, HIB19, SJ25C1, J3-119. Plots show the MFI (geometric mean of fluorescence intensity) from anti-CD19 flow cytometry staining of tumor cells. Data are presented as the mean ± SD (I, II, V, VII, VIII) or median with IQR (III, IV, VI) from 5-16 biological replicates. Statistical analysis was performed using the Welch F test (I, II, V, VII) or one-way ANOVA (VIII) with Tukey's post-hoc test or Kruskal-Wallis test (III, IV, VI) with Dunn's post-hoc test for multiple comparisons.

**Supplementary Figure 2**

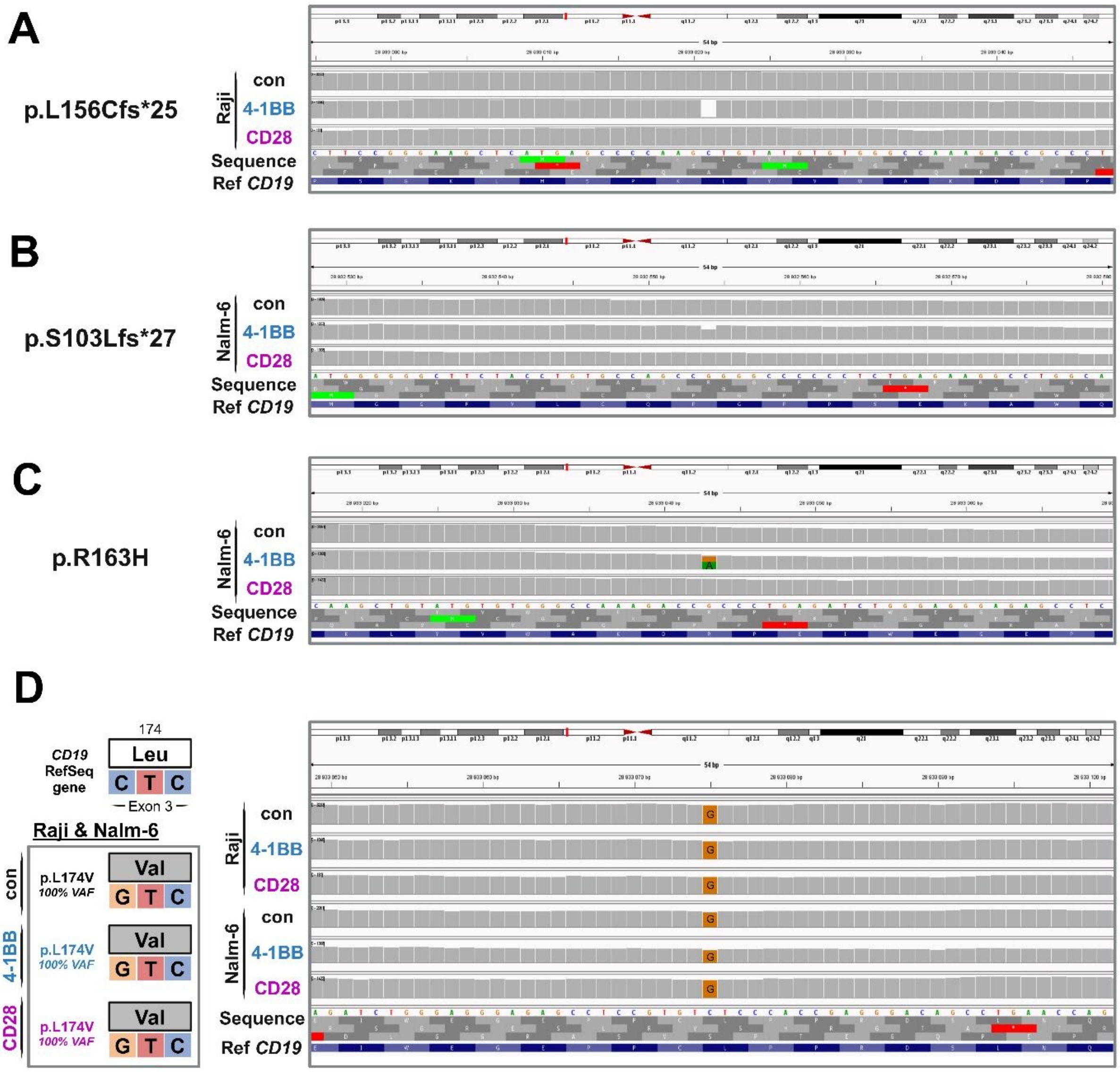


**Supplementary Figure 2.**

(**A-C**) IGV's screenshots of *CD19* sequences from Raji/Nalm-6 control (con), CAR-4-1BB and CAR-CD28 cell lines showing particular mutations p.S103Lfs*27 (**A**), p.L156Cfs*25 (**B**) and p.R163L (**C**). The CD19 sequences were visualized with IGV based on pooled RNAseq data from 3 biological replicates of each variant.

(**D**) The scheme of *CD19* sequence in Raji and Nalm-6 control (con), CAR-4-1BB and CAR-CD28 detected in RNAseq data showing the polymorphism p.L174V (left panel). An IGV's screenshot of *CD19* sequences showing the same polymorphism (right panel).

### Supplementary Figure 3

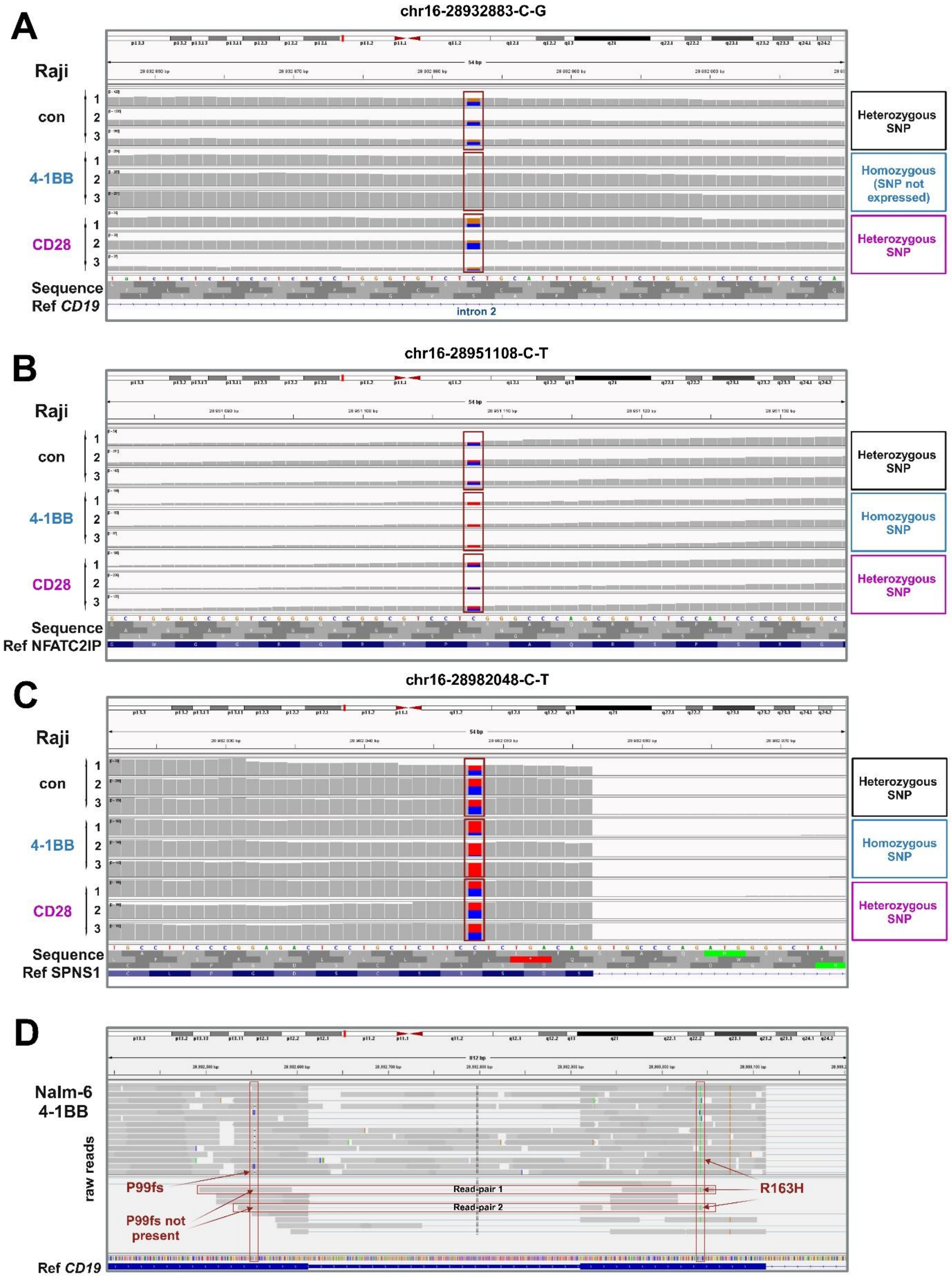

**Supplementary Figure 3.**

(**A-C**) IGV's screenshots showing parts of *CD19 (A), NFATC2IP (B) and SPNS1* gene sequences from Raji control (con), CAR-4-1BB and CAR-CD28 cell lines with the particular SNPs. Red frames highlighted the point mutations. The sequences were visualized with IGV based on RNAseq data from 3 biological replicates of each variant.

(**D**) The IGV screenshot of raw reads data from the Nalm-6 CAR-4-1BB cell line. The red frames show 2 read-pairs covering both mutation sites detected in Nalm6 CAR-4-1BB variant (p.R163H) and deletion in codon P99 resulting in frameshift mutation (p.S103Lfs*27). The data comes from one sample of 3 biological replicates.

**Supplementary Figure 4**

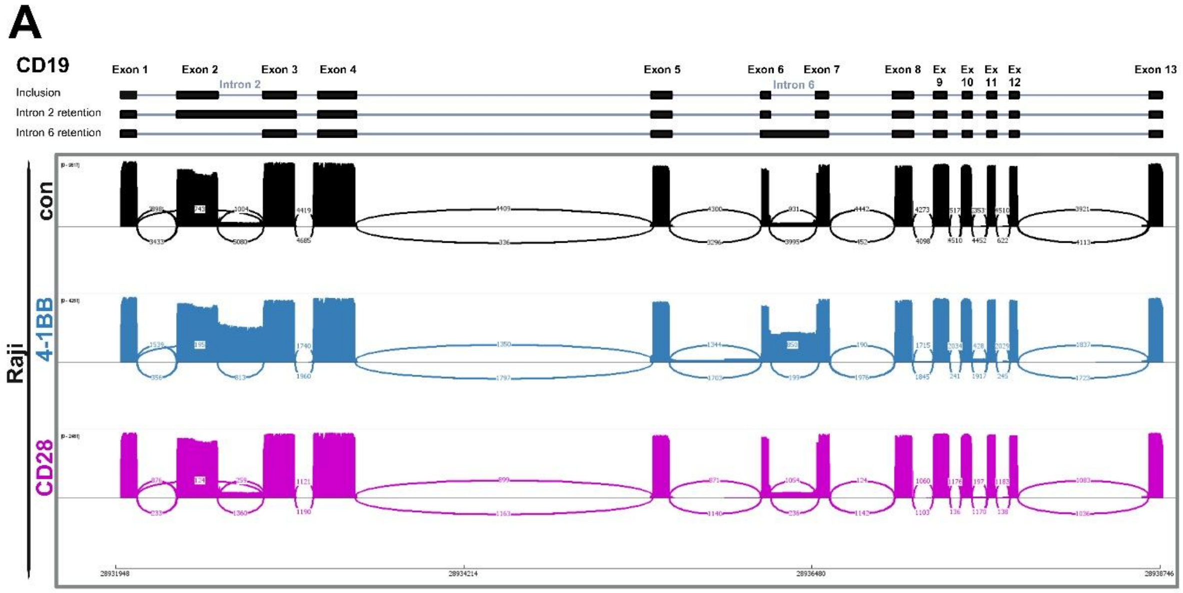


**Supplementary Figure 4.**

(**A**) Sashimi plots with coverage and the number of junction reads in Raji control (black), 4-1BB (blue) and CD28 (magenta) showing the intron 2 and intron 6 retention. Plots show the data from long-read (lr) sequencing of *CD19* amplicone using Oxford Nanopore Technology.

**Supplementary Figure 5**

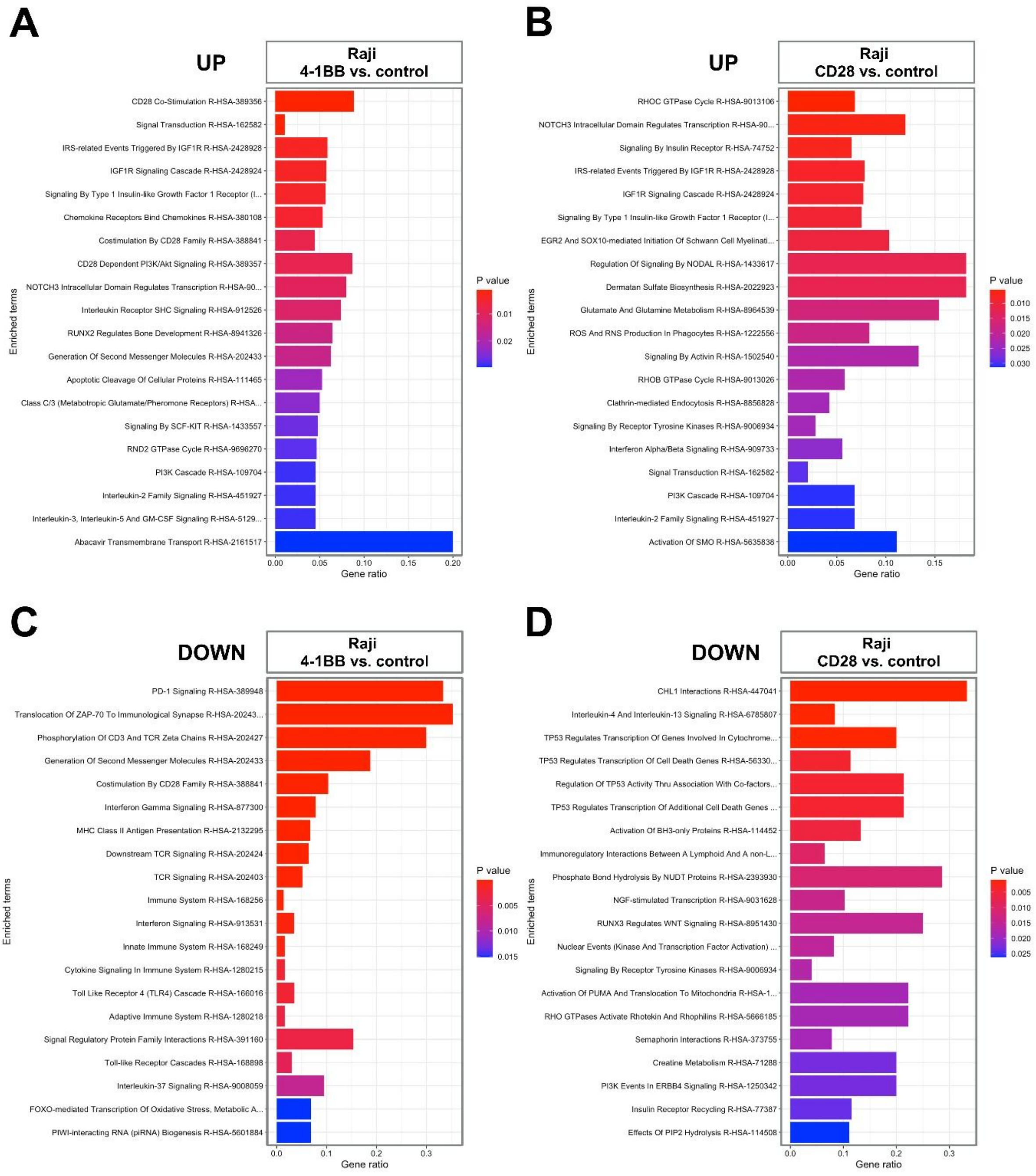


**Supplementary Figure 5.**

**(A-D)** The analysis of the significantly changed signaling pathways in Raji cells based on Reactome database. The charts show enriched upregulated (UP) pathways between control and CAR-4-1BB (**A**) or CAR-CD28 (**B**), and significantly enriched downregulated (DOWN) pathways between control and CAR-4-1BB (**C**) or CAR-CD28 (**D**).

**Supplementary Figure 6**

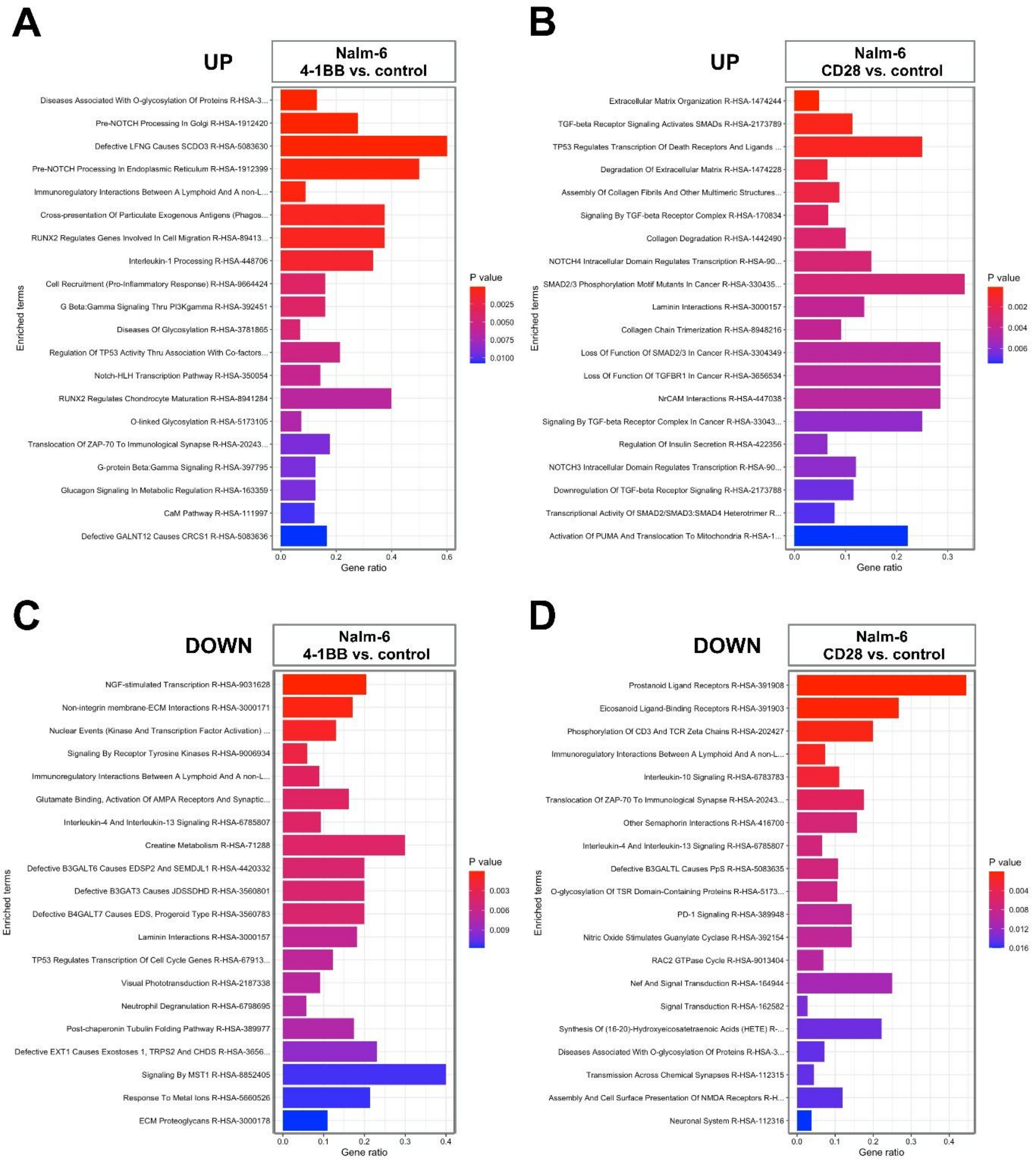


**Supplementary Figure 6.**

**(A-D)** The analysis of the significantly changed signaling pathways in Nalm-6 cells based on Reactome database. The charts show enriched upregulated (UP) pathways between control and CAR-4-1BB (**A**) or CAR-CD28 (**B**), and significantly enriched downregulated (DOWN) pathways between control and CAR-4-1BB (**C**) or CAR-CD28 (**D**).

**Supplementary Figure 7**

**A**

| parameters used in simulation | CD19-4-1BB CAR-T cells | CD19-CD28 CAR-T cells |
|---|---|---|
| initial number of resistant cells | 1 | 1 |
| initial number of low antigen cells | 9089 | 9089 |
| initial number of high antigen cells | 9986480 | 9986480 |
| resistant cells antigen max value | 10 | 10 |
| low antigen max value | 1000 | 1000 |
| high antigen min value | 1000 | 1000 |
| mutation probability | 0.00005 | 0.00005 |
| high antigen cell death probability | 0.1 | 0.1 |
| low antigen cell death probability | 0.02 | 0.1 |

**B**

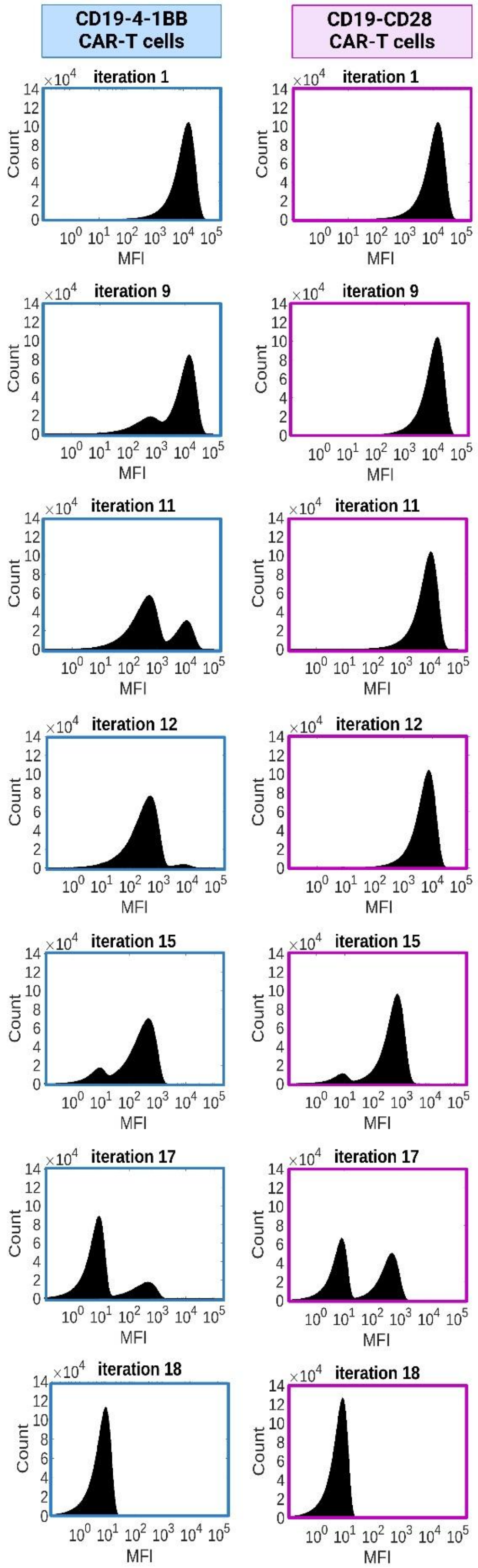


**Supplementary Figure 7.**

**(A)** Main parameters set for simulations.

**(B)** Histograms showing changes in CD19 MFI in subsequent passages (iterations) mimicking contact with CD19-4-1BB CAR-T cells (left) or CD19-CD28 CAR-T cells (right). The graphs show selected iteration from 19 that were performed.
